# Optimization of a bacterial three-hybrid assay through *in vivo* titration of an RNA-DNA adapter-protein

**DOI:** 10.1101/2020.07.23.216291

**Authors:** Clara D. Wang, Rachel Mansky, Hannah LeBlanc, Chandra M. Gravel, Katherine E. Berry

## Abstract

Non-coding RNAs regulate gene expression in every domain of life. In bacteria, small RNAs (sRNAs) regulate gene expression in response to stress and are often assisted by RNA-chaperone proteins, such as Hfq. We have recently developed a bacterial three-hybrid (B3H) assay that detects the strong binding interactions of certain *E. coli* sRNAs with proteins Hfq and ProQ. Despite the promise of this system, the signal-to-noise has made it challenging to detect weaker interactions. In this work, we use Hfq-sRNA interactions as a model system to optimize the B3H assay, so that weaker RNA-protein interactions can be more reliably detected. We find that the concentration of the RNA-DNA adapter is an important parameter in determining the signal in the system, and have modified the plasmid expressing this component to tune its concentration to optimal levels. In addition, we have systematically perturbed the binding affinity of Hfq-RNA interactions to define, for the first time, the relationship between B3H signal and *in vitro* binding energetics. The new pAdapter construct presented here substantially expands the range of detectable interactions in the B3H assay, broadening its utility. This improved assay will increase the likelihood of identifying novel protein-RNA interactions with the B3H system, and will facilitate exploration of the binding mechanisms of these interactions.

## NTRODUCTION

Bacteria are capable of exquisite control of gene expression in response to cellular or environmental stress.(Gottesman and Storz 2011; Wagner and Romby 2015; Dersch et al. 2017) Small RNAs (sRNAs) are a class of *trans*-acting non-coding RNAs in bacteria that are expressed under specific conditions to regulate groups of messenger RNAs (mRNAs).(Gottesman and Storz 2011; Wagner and Romby 2015; Hör et al. 2020; Quendera et al. 2020) Many bacterial sRNAs are known to depend on a protein cofactor, such as the RNA-chaperone protein Hfq, for their function.(Holmqvist and Vogel 2018; Updegrove et al. 2016; Gottesman and Storz 2015; Vogel and Luisi 2011; Chao and Vogel 2010; Zhang et al. 2003; Schu et al. 2015) Indeed, RNA-protein interactions involving both coding and non-coding RNAs underlie many aspects of post-transcriptional gene regulation in the bacterial stress response.

Despite these important roles, the full spectrum of proteins that interact with bacterial regulatory RNAs has not yet been defined, and many RNA-binding proteins still have unclear mechanisms of action. Numerous methods have therefore been developed to identify and study RNA-protein interactions, ranging from co-immunoprecipitation (Andresen and Holmqvist 2018; Saulière and Le Hir 2015) and co-sedimentation (Smirnov et al. 2016) techniques to biochemical assays that provide access to quantitative, physical parameters such as binding energies and rates (Faner and Feig 2013; Glover et al. 2015). In addition, a set of molecular-genetic assays have been developed to allow RNA-protein interactions to be studied in a cellular environment.(Koh and Wickens 2014a, 2014b; Bernstein et al. 2002; Gelderman et al. 2015; SenGupta et al. 1996) Such genetic assays provide a powerful complement to biochemical methods and provide pathways to use both forward and reverse genetic approaches to probe RNA-protein interactions.

We have recently developed a bacterial three-hybrid (B3H) assay that connects the strength of sRNA-protein interactions inside of living *E. coli* cells to the output of a reporter gene.(Berry and Hochschild 2018; Pandey et al. 2020) In this three-hybrid system, expression of two hybrid proteins and one hybrid RNA link the transcription of a β-galactosidase (β-gal) gene to the strength of an RNA-protein interaction of interest. An RNA-DNA adapter protein (hereafter, “adapter” or CI-MS2^CP^) tethers the hybrid “bait” RNA upstream of a weak core promoter, where it can interact with an RNA polymerase (RNAP)-bound “prey” protein (Fig. 1A). While certain sRNA-Hfq interactions are well detected in this assay,(Berry and Hochschild 2018) the signal-to-noise for many well-characterized protein-RNA interactions is not strong enough for interactions to be reliably detected. In addition, there has been no systematic examination of the relationship between the signal from the B3H assay and the binding energy of a given RNA-protein interaction. Increased signal would substantially expand the utility of this assay for a wider range of RNA-binding proteins.

**Figure 1.**
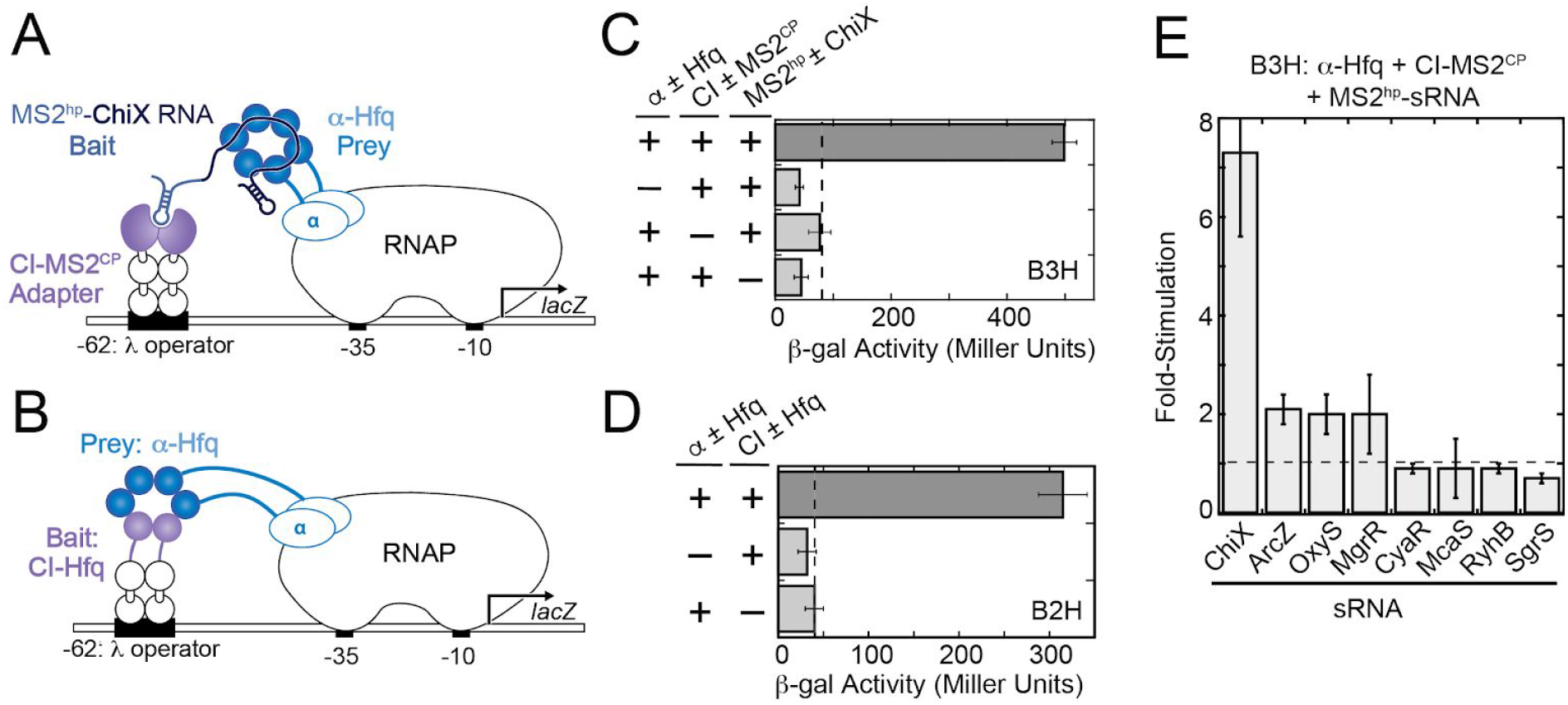
Bacterial two- and three-hybrid interactions with Hfq. (**A**) Schematic showing B3H system for detection of Hfq-RNA interactions. Interaction between an Hfq “prey” protein moiety and RNA “bait” moiety activates transcription from a test promoter, which directs transcription of a *lacZ* reporter gene. The test promoter (p*lac*-O_L_2–62), bears the λ operator O_L_2 centered at position –62 relative to the transcription start site. The RNA-binding moiety MS2^CP^ is fused to λCI (CI-MS2^CP^) as an RNA-DNA adapter (“Adapter”) to tether the hybrid RNA (MS2^hp^-ChiX) to the test promoter. (**B**) Schematic showing B2H system. Interaction between Hfq protein moieties fused, respectively, to α-NTD (α-Hfq) as “Prey” and to λCI protein (CI) as “Bait” activates transcription of *lacZ* from the same O_L_2–62 test promoter as above. (**C**) Results of β-galactosidase (β-gal) assays performed in KB473 cells transformed with three compatible plasmids: pPrey (α±Hfq) that encoded α alone (–) or the α-Hfq (+) fusion protein, pAdapter (CI ± MS2^CP^) that encoded λCI alone (–) or the CI-MS2^CP^ fusion protein (+), and a pBait (MS2^hp^ ± ChiX) that encoded a hybrid RNA with the ChiX sequence following two copies of an MS2^hp^ moiety (+) or an RNA that contained only the MS2^hp^ moieties (–). **(D)** Results of β-gal assays performed in KB473 containing three compatible plasmids: one (α±Hfq that encoded α alone (–) or the α-Hfq (+) fusion protein and another (CI±Hfq) that encoded λCI alone (–) or the CI-Hfq fusion protein (+). **(E)** Results of B3H β-gal assays performed in KB473 cells with pAdapter (CI-MS2^CP^, pKB989), pPrey (α-Hfq, pKB817) and pBaitRNA (2xMS2^hp^ moieties fused to the listed sRNAs). The corresponding negative controls as in (C) were also conducted in parallel. The bar graph here, and for all subsequent B2H and B3H experiments, shows the fold-stimulation over basal levels: that is, the β-gal activity measured in the presence of all hybrid constructs divided by the activity of the highest negative control sample expressing only half of one hybrid construct. The dashed horizontal line represents “basal” levels. Data are presented as the average value of three independent measurements and error bars show the standard deviation of these values.

Here we present our efforts to optimize expression of the adapter protein, as a strategy to increase overall assay signal and sensitivity. We have performed an *in vivo* titration of this key component in order to understand the relationship between its concentration and B3H signal and have utilized both forward and reverse genetic approaches to improve the signal of the assay. We introduce an optimized adapter-encoding plasmid (pAdapter), and demonstrate that it improves B3H signal for multiple sRNA-Hfq interactions. In addition, we have used a panel of Hfq variants with established effects on the energetics of *in vitro* RNA binding, in order to define the relationship between *in vitro* binding affinity and *in vivo* B3H signal. This work provides a more quantitative benchmark for interpreting data from this system and expands the useful applications of this assay.

## METHODS

Bacterial strains, plasmids and oligonucleotides used in this study are specified in Tables 1-3. NEB5α was the recipient strain for all cloned plasmids and was purchased from New England Biolabs. Each strain and plasmid confers resistance to an antibiotic, abbreviated as follows: tetracycline (tet), kanamycin (kan), carbenicillin (carb; an analog of ampicillin, amp) chloramphenicol (cm), and spectinomycin (spec).

### Plasmid construction and mutagenesis

Plasmids were constructed as specified in Table S1 and the construction of key parent vectors is described below. Plasmids pCW13-19 (pAC-p_constit_-λCI-MS2^CP^) were constructed by mutagenic PCR from pCW17.(Pandey et al. 2020) PCR mutagenesis to create site-directed mutants of the promoter region of pCW17 or the *hfq* sequence in pKB817 was conducted either with Quikchange-style mutagenesis or with Q5 Site-Directed Mutagenesis (New England Biolabs) using end-to-end primers designed with NEBaseChanger.(Table S2 and S3)

All 2xMS2^hp^–sRNA hybrids were constructed by inserting the sRNA of interest into the XmaI/HindIII sites of pKB845 (pCDF–pBAD–2xMS2^hp^–XmaI–HindIII).(Berry and Hochschild 2018) pHL34 (pCDF-pBAD-2xMS2^hp^-ΔXmaI-DsrA) was further derived from pKB941 (pCDF-pBAD-2xMS2^hp^-DsrA) using Q5 mutagenesis to remove the XmaI restriction site (6bp; Table S2). No terminator sequence outside of the intrinsic terminators in each sRNA was provided, except for a *trpA* terminator present in pHL26 (pCDF-pBAD-1xMS2^hp^-A_27_-T_trpA_); this construct was derived through insertion of a poly(adenosine) stretch into pHL6 (pCDF-pBAD-1xMS2^hp^-T_trpA_ Pandey et al. 2020), using Q5 Site-Directed Mutagenesis (Table S2).

Plasmid libraries introducing random mutations into the −35-promoter region and Shine-Dalgarno region of pAdapter (−35 and SD libraries, respectively) were generated from pCW17 by 25 rounds of PCR amplification of the promoter region using mutagenic PCR with primers oCW36 and oCW37 (−35 library) and oCW40 and oCW41 (SD library). These primers possessed regions of complementarity to pCW17 surrounding a randomized hexameric sequence at either the −35 hexamer or the center of the Shine-Dalgarno sequence. Within this NNNNNN sequence, each position had a 25% possibility of each base. Following KLD (kinase, ligase, and DpnI) treatment and transformation into NEB 5-alpha F’Iq cells (New England Biolabs), cells were grown as near-lawns on LB-chloramphenicol plates and a miniprep was performed from a resuspension of ~5000 colonies for each library.

### β-galactosidase assays

For B3H assays, two types of reporter cells were used (KB473 or KB483).(Berry and Hochschild 2018; Pandey et al. 2020) Both strains feature the same p*lac*-O_L_2–62-*lacZ* reporter and have *hfq* deleted from their genome. While the F’ episome in KB473 carrying the p*lac*-O_L_2–62-*lacZ* test promoter confers kanamycin resistance, the corresponding F’ episome in KB483 confers tetracycline resistance. For B3H assays using either reporter strain, reporter cells were transformed with three plasmids: one expressing a CI-MS2^CP^ adapter protein (pAdapter), another expressing α-Hfq (pPrey), and a third expressing a 2XMS2^hp^-RNA (pBaitRNA), or a corresponding plasmid expressing half of each hybrid construct (α alone, CI alone or the MS2^hp^ moieties alone alone).

For liquid assays, three colonies from each transformation were picked and inoculated into 1 mL LB broth containing cm (25μg/mL), carb (100μg/mL), spec (100μg/mL) with either kan (50μg/mL; for KB473 cells) or tet (10μg/mL; for KB483 cells), and 0.2% arabinose in a 96-well deep well plate (Greiner Bio-One). The deep-well plates were covered with a breathable film (VWR) and shaken at 900 rpm in a shaker (Benchmark Scientific) at 37°C overnight. The following day, the overnight cultures were back-diluted 1:50 in 200 μL LB supplemented with the same antibiotics and inducers as above. Cells were grown to mid-log (OD_600_=0.4-0.6) in covered 96-well plates in a shaker at 37°C. Lysis of mid-log cells and measurement of β-gal activity was performed as previously described.(Berry and Hochschild 2018) Isopropyl-β-D-thiogalactoside (IPTG) was added as indicated in figure legends. Unless otherwise noted, B3H assays were conducted in KB483 cells grown in the absence of IPTG and the presence of 0.2% arabinose. When present in both overnight cultures and back-dilutions, IPTG concentration is indicated with an asterisk (*); otherwise IPTG was absent from overnight cultures and only present in back-dilutions.

For plate-based assays, cells were transformed and grown to mid-log as above. Mid-log cells (2 μL after 1:100 dilution) were plated on X-gal-indicator medium (containing 40μg/mL 5-bromo-4-chloro-3-indolyl-β-D-galactopyranoside (X-gal), 125 μM phenylethyl-β- D-thiogalactopyranoside (TPEG), and 1.5μM IPTG, along with arabinose and antibiotics as above) and incubated overnight. Following overnight growth, plates were allowed to rest at 4°C for 1-2 days to allow color to develop, and were photographed with a black-velvet background with oblique lighting. The brightness and levels of photographs were adjusted evenly across images.

For both liquid and plate-based assays, B3H interactions are interpreted as the β-gal activity in reporter cells containing all hybrid constructs (α-Hfq, CI-MS2^CP^ and MS2^hp^-RNA), relative to negative controls. For qualitative plate-based assays, this is represented as the relative color of bacterial patches on X-gal containing media, where blue color represents higher β-gal activity, and therefore stronger RNA-protein interaction. For liquid assays, except in Fig. 1C, this is reported as the fold-stimulation over basal levels; this is the β-gal activity (expressed in Miller units) in reporter cells containing all hybrid constructs divided by the highest activity from negative controls in which one of the full hybrid constructs was replaced by a plasmid expressing half of each construct (α alone, CI alone or the MS2^hp^ moieties alone alone).

B2H assays were conducted as above with the following modifications: spectinomycin and arabinose were left out of liquid and solid media and only two plasmids were transformed into cells: one expressing the CI-Bait (*i.e*. CI-Hfq, CI-MS2^CP^ or CI-β-flap), another expressing α-Prey (*i.e*. α-Hfq, α-MS2^CP^, or pBR-σ^70^) or a corresponding plasmid expressing half of each hybrid construct (α alone or CI alone). B2H interactions, except in Fig. 1D, are reported as fold-stimulation over basal levels, as described above.

For CI-repression assays, reporter-strain cells (FW123) were transformed with plasmids containing the pAC backbone with CI or a negative control plasmid containing the pAC backbone without the CI protein (ΔCI).(Pinkett et al. 2006; Whipple et al. 1994) Three colonies from each transformation were inoculated into 1 mL LB containing cm (25 μg/mL), kan (50 μg/mL) and IPTG as indicated in a 96-well deep well plate. Cells were grown in this medium overnight, grown to mid-log, lysed and assayed for β-gal activity as above.

All assays were conducted in biological triplicate and repeated independently on at least three days, and a representative data set is shown. Data are plotted as the average across three triplicate measurements and error bars represent one standard deviation above and below the mean.

### Forward genetic screen

Mutant plasmid libraries were transformed into KB473 cells already carrying pKB817 (α-Hfq) and pKB856 (pCDF-2xMS2^hp^-MgrR). Cells were grown on LB agar containing the antibiotics cm (25μg/mL), carb (100μg/mL), kan (50μg/mL), spec (100μg/mL), inducers IPTG (1.5μM) and arabinose (0.2%), and indicators X-gal (40μg/mL) and TPEG (125μM). Plates were grown overnight at 37°C and stored at 4°C for an additional 24 hours. Colonies that appeared dark blue were restreaked to confirm colony color on the same plate conditions as above. Plasmids from colonies confirmed to be darker blue than starting colonies were miniprepped to be tested in liquid β-gal assays, as above. Promising candidates were analyzed by DNA sequencing with primers oCW5 and oRM20.

## RESULTS

### Introduction of Bacterial Two- and Three-Hybrid Assay

Our B3H system and the B2H system from which it was adapted both depend on ‘bait’ and ‘prey’ moieties expressed as hybrid molecules tethered to the *E. coli* transcriptional machinery. In this configuration, interaction between bait and prey molecules activates a reporter gene (*lacZ*; Fig 1A,B).(Dove et al. 1997; Dove and Hochschild 2001; Nickels et al. 2002; Berry and Hochschild 2018) Both systems tether prey proteins to RNAP via fusion to the N-terminal domain of the alpha subunit of RNAP (*e.g.* α-Hfq) and present bait molecules via CI-fusion proteins bound at a λ-operator (O_L_2) sequence upstream of the test promoter. However, the B2H and B3H systems differ in whether the bait molecule is protein or RNA. In the B3H system, a CI-MS2^CP^ fusion protein serves as an RNA-DNA adapter protein (“adapter”) and recruits an MS2^hp^-containing RNA as bait (*e.g.* MS2^hp^-ChiX; Fig. 1A). In the B2H system, a CI-fusion protein (*e.g.* CI-Hfq) directly presents the protein bait (Fig. 1B). The physical interaction between bait and prey moieties stabilizes RNAP at the p*lac*-O_L_2–62 test promoter and stimulates *lacZ* transcription.

As previously reported, a B3H interaction between Hfq and ChiX results in robust stimulation of *lacZ* transcription above the basal levels detected when half of any hybrid component is omitted (Fig. 1C).(Berry and Hochschild 2018) Similarly, an Hfq-Hfq B2H interaction results in strong stimulation of β-gal activity above basal levels (Fig 1D). While many B2H interactions produce similar signal as an Hfq-Hfq B2H interaction,(Dove and Hochschild 2001) the ChiX-Hfq B3H interaction produces uniquely high signal among Hfq-sRNA B3H interactions.(Berry and Hochschild 2018) Indeed, several other well-established Hfq-dependent sRNAs produce much lower signal than that observed for a ChiX-Hfq interaction (Fig. 1E).

### IPTG-dependence of B2H and B3H signal

In both the B2H and B3H system, each hybrid component is provided on a separate compatible plasmid (Fig. 2A).(Berry and Hochschild 2018) In the B2H system, both hybrid proteins are expressed from IPTG-inducible promoters from plasmids we call “pPrey” and “pBait”. In the original iteration of the B3H system, the pAC-λCI vector (pAdapter) expressed CI-MS2^CP^ in an IPTG-dependent manner, as it was created directly from the IPTG-inducible plasmid expressing CI-bait for B2H assays. The MS2^hp^-hybrid RNA in the B3H system is provided on a third compatible plasmid (pBait_RNA_) and is induced by arabinose.(Berry and Hochschild 2018)

**Figure 2.**
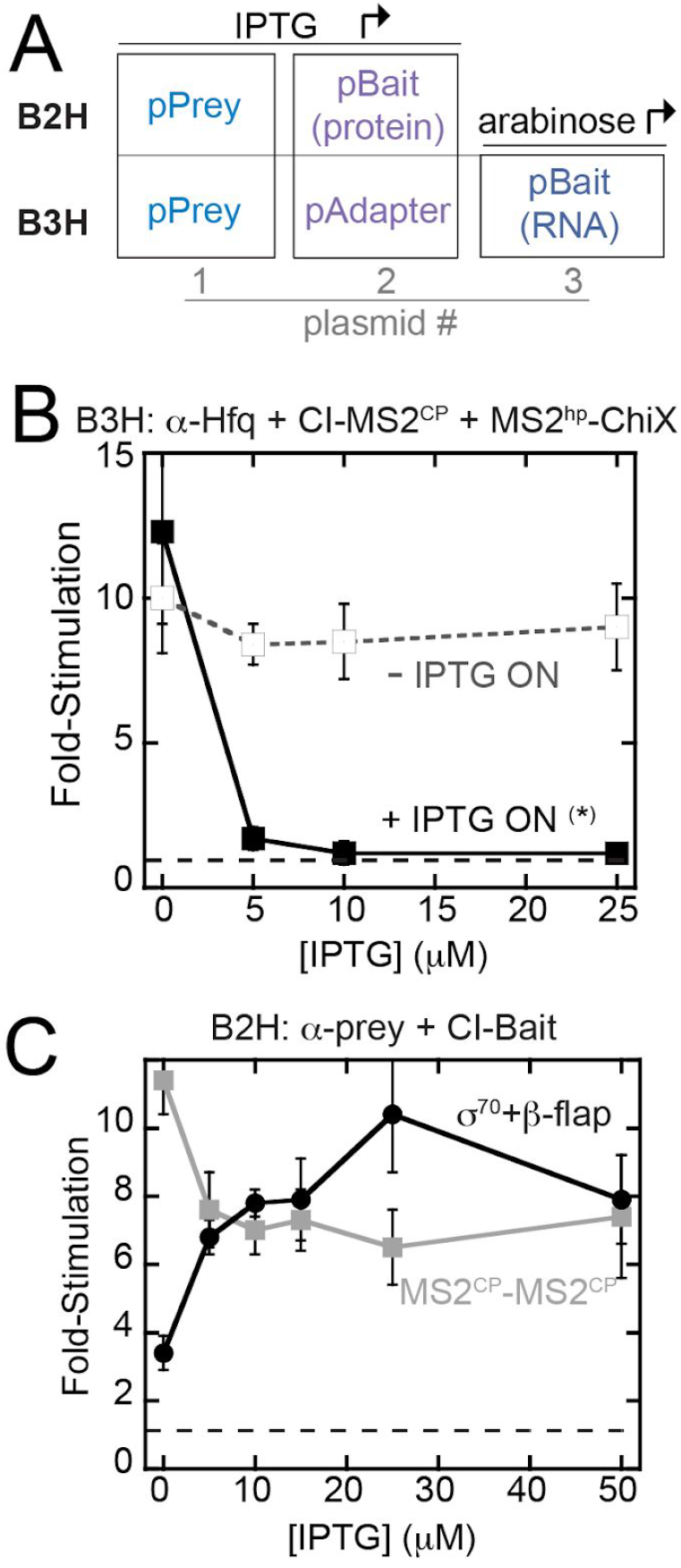
Bacterial two- and three-hybrid interactions display distinct patterns of IPTG-dependence. (A) Schematic showing the plasmids used to express bait and prey hybrid components in two-hybrid (B2H) and three-hybrid (B3H) assays and, in the case of the B3H assay, the adapter protein. Inducers that drive expression of each component are indicated. Results of β-gal assays detecting (B) B3H or (C) B2H assays when reporter cells were grown in the presence of varying IPTG concentrations. (B) Results of B3H β-gal assays performed in KB473 cells with pAdapter (CI-MS2^CP^, pKB989), pPrey (α-Hfq) and pBaitRNA (2xMS2^hp^-ChiX, pKB909). Reporter cells were grown under two conditions: one in which IPTG was added at the indicated concentration in both the overnight and back-dilution cultures (+ IPTG ON *) and another in which IPTG was added only in the back-dilution cultures (- IPTG ON). (C) Results of B2H β-gal assays performed in KB483 cells with pBait (MS2^CP^: pKB989 or β-flap: pACλCI-β-flap^831-1057^, pPrey (MS2^CP^: pKB815 or σ^70^: pBRa-σ70^D581G^), grown at the indicated concentration of IPTG.

In an effort to optimize assay conditions and increase signal-to-noise in the B3H assay, we began by investigating the dependence of B3H signal on the levels of the protein components. To do so, we compared the strength of a ChiX-Hfq B3H interaction as a function of IPTG concentration. Surprisingly, we found that B3H signal decreased as IPTG concentration increased (Fig. 2B). This negative IPTG-dependence of B3H signal was especially apparent when IPTG was present in overnight cultures (+ IPTG ON; indicated with *). This result suggested that at least one of the fusion proteins was inhibitory to B3H signal at higher concentrations. Recall that in these original B3H vectors used, both the RNA-DNA adapter and the prey-fusion protein were induced by IPTG (Fig. 2A).

To explore the basis for the counterintuitive dependence on protein expression, we compared the IPTG-dependence of the B3H system to that of simpler B2H assays. We compared two protein-protein interactions in the B2H system: one between a region of the RNAP β-flap previously shown to strongly interact with a σ^70^ mutant in this system,(Dove and Hochschild 2001) and one between two copies of the MS2^CP^ used as the adapter protein in B3H assays. (Berry and Hochschild 2018) As previously observed, a B2H interaction between σ70^D581G^ and β-flap^831-1057^ produces β-gal signal with a positive dependence on IPTG;(Dove and Hochschild 2001) the more fusion protein that is expressed, the higher the fold-stimulation of *lacZ* expression (Fig 2C). Interestingly, and in contrast, the B2H interaction between CI-MS2^CP^ (the RNA-DNA adapter used in the B3H system) and α-MS2^CP^ shows a modest negative dependence on IPTG concentration: the strongest interaction is seen when both fusion proteins are uninduced.

We reasoned that the negative-IPTG dependence in the B3H system could be due to an inhibitory effect of either the prey or adapter component, as both were originally induced by IPTG (Fig 2A). Based on the negative IPTG-dependence of the MS2^CP^-MS2^CP^ B2H interaction, we favored the possibility that high levels of CI-MS2^CP^, encoded by pAdapter, inhibited B3H signal. We further reasoned that, since MS2^CP^ is a viral coat-protein that evolved to form higher-order structures, such structures could be problematic inside of the cell (Fig. S1A). In addition, the λ-operator site is present as a single-copy on an episomally encoded reporter whereas α-Hfq can likely bind thousands of copies of RNAP in the cell. This difference in binding-site abundance could mean that, while very little CI-MS2^CP^ adapter is needed to saturate the λ-operator DNA, too much could result in capping interactions of the DNA and hybrid RNA by separate adapter molecules (Fig. S1B). Such capping interactions would remove the stabilizing effect of an RNA-protein interaction at the test promoter.

### Development of a constitutive pAdapter

If high levels of CI-MS2^CP^ adapter protein inhibit B3H signal, we reasoned it would be helpful for the concentration of prey-fusion protein to be adjustable independently from that of the adapter. We therefore removed the *lacO* site from the *lacUV5* promoter in the original pAdapter construct (pKB989) to create a version of this plasmid to generate constitutive expression of CI-MS2^CP^ (pCW18; Fig. 3A). At the same time as removing the *lacO* site, additional mismatches relative to the consensus σ^70^ promoter-sequence were introduced to the −10 and −35 regions of pCW18’s promoter to weaken the expression of adapter (Fig. 3A).

**Figure 3.**
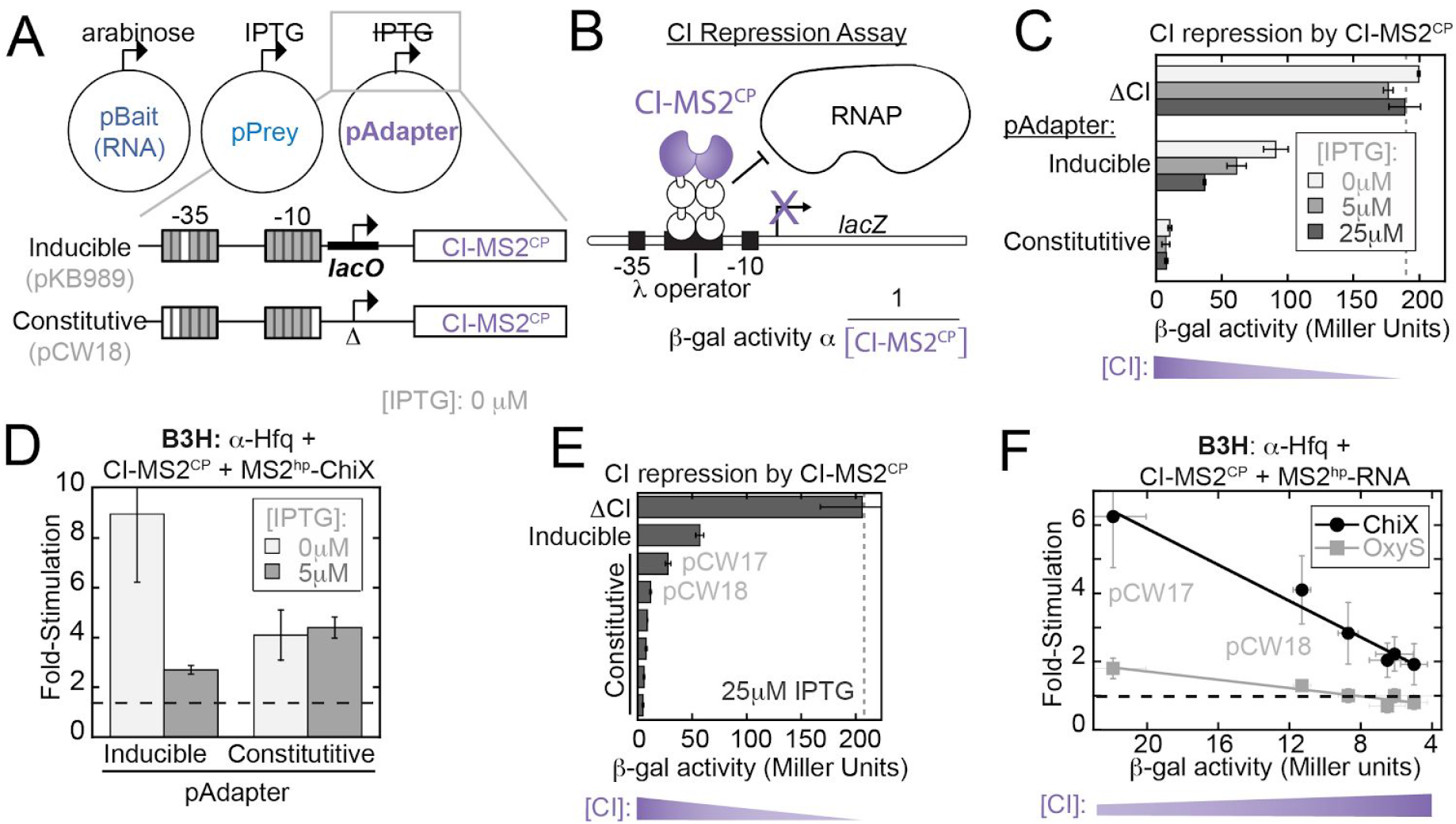
Constitutive expression of the RNA-DNA adapter eliminates negative IPTG-dependence of the B3H assay. (A) Schematic of B3H plasmid constructs along with inducers expressing each hybrid component. Inset shows changes to the pAdapter promoter sequence to eliminate IPTG-inducible expression of CI-MS2^CP^ and create pCW18 as a constitutive pAdapter construct. Grey boxes represent matches to and white boxes mismatches from the consensus sequence for σ^70^ promoters. See Table S4 for details. (B) Schematic showing the logic of the CI-repression assay using FW123 reporter cells (Pinkett et al. 2006; Whipple et al. 1994) A single-copy test promoter driving *lacZ* expression bears a λ operator between the −35 and −10 promoter elements. Binding of CI repressor at this site represses *lacZ* transcription. The relative expression of a CI fusion protein is inversely proportional (α) to β-gal activity. (C) Results of CI-repression β-gal assays. FW123 cells carrying a single pAdapter plasmid expressing either the CI-MS2^CP^ fusion driven by an IPTG-Inducible promoter (Inducible; pKB989) or a constitutive promoter (constitutive; pCW18) were grown at the indicated concentration of IPTG. β-gal values were compared to baseline levels set by a corresponding plasmid lacking any CI fusion protein (ΔCI). (D) Results of B3H β-gal assays performed in KB473 cells with pAdapter (CI-MS2^CP^ fusion expressed from pKB989 (inducible) or pCW18 (constitutive), pPrey (α-Hfq) and pBaitRNA (2xMS2^hp^-ChiX). (E) Results of CI-repression β-gal assays from FW123 cells containing ΔCI, pKB989 (Inducible), or, from top to bottom, pCW17,18,19,14,15,16 (constitutive), grown in the presence of 25 μM IPTG. (F) Results of B3H β-gal assays performed in KB473 cells, with pPrey (α-Hfq), pBait (2xMS2^hp^-ChiX or 2xMS2^hp^-OxyS, pKB912) and pAdapter (from left to right: pCW17,18,19,16,14,15). B3H results are plotted against values for each pAdapter in CI repression assays, collected as in (E) but with cells grown in the absence of IPTG.

In order to compare expression levels of CI-MS2^CP^ across different pAdapter constructs, we used a CI-repression assay. In this assay, reporter cells (FW123) carry a single-copy test promoter bearing the λ operator between the −35 and −10 regions of the *lacZ* promoter (Fig 3B).(Pinkett et al. 2006; Whipple et al. 1994) In this configuration, the CI protein acts as a transcriptional repressor: binding of CI at the λ-operator represses *lacZ* transcription by physically blocking RNAP (Fig. 3B). β-gal activity in FW123 reporter cells is compared to a baseline of unrepressed transcriptional activity, determined by *lacZ* expression observed in cells with a plasmid lacking the CI protein entirely (ΔCI). The relative expression of a CI-fusion protein, such as CI-MS2^CP^, can be inferred from an inverse relationship with β-gal activity.

We began by comparing CI-MS2^CP^ expression, as a function of IPTG concentration, from pAdapter constructs bearing either the inducible or constitutive promoter. Cells carrying the inducible pAdapter construct demonstrate an IPTG-dependent increase in CI-MS2^CP^ levels, whereas *lacZ* expression remains essentially unchanged in the presence of IPTG when cells carry the constitutive pAdapter (pCW18; Fig. 3C). It is interesting to note that adapter expression from pCW18, while independent of IPTG concentration, was stronger than we expected based on the design of the promoter sequence (see below). Nevertheless, these results confirmed that removal of the *lacO* site from pKB989 indeed resulted in a constitutive pAdapter construct, pCW18.

### Constitutive adapter expression eliminates IPTG-inhibition of B3H signal

To test whether constitutive expression of CI-MS2^CP^ eliminates the negative relationship between B3H signal and fusion-protein induction by IPTG, we compared Hfq-ChiX B3H interactions using either the inducible or constitutive pAdapter constructs. As previously observed, reporter cells carrying plasmids for the Hfq-ChiX interaction along with an IPTG-inducible pAdapter (pKB989) show a dramatic decrease in fold-stimulation when IPTG is present (Fig. 3D; left). However, when CI-MS2^CP^ was expressed from a constitutive promoter (pCW18), B3H signal was essentially unchanged upon addition of IPTG (Fig. 3D). When the B3H assay is conducted with this constitutive pAdapter construct, only the prey protein (α-Hfq) is induced by IPTG. Thus, this result is consistent with our hypothesis that observed inhibition of B3H signal by IPTG was driven by high levels of the CI-MS2^CP^ adapter protein. While use of a constitutive pAdapter eliminated negative dependence of assay signal on IPTG, it still yielded lower signal than the inducible pAdapter in the absence of IPTG (Fig. 3D). This suggested that further tuning of the constitutive level of CI-MS2^CP^ expression would be necessary for optimal signal.

### Tuning adapter levels through varied constitutive promoters

In order to probe the effects of a wider range of CI-MS2^CP^ concentrations, we created a set of pAdapter constructs with constitutive promoters of varying strength. Mutations were made to both the −35 and −10 promoter elements to adjust predicted promoter strength (Table S4). Adapter expression from each construct was then assessed using the CI-repression assay. The range of CI-MS2^CP^ levels produced by this panel (Fig. 3E) afforded the opportunity to systematically investigate dependence of B3H signal across a range of adapter concentrations. To do so, we measured B3H signal for Hfq interactions with ChiX and OxyS sRNAs, using this panel of pAdapter constructs. The results demonstrated a striking inverse relationship between the B3H signal generated by each pAdapter construct and the amount of CI-MS2^CP^ it produces (Fig. 3F). Of this set of pAdapter constructs, pCW17 displayed the highest fold-stimulation in the B3H assay, resulting in approximately 6- and 2-fold stimulation over basal levels for ChiX and OxyS interactions with Hfq, respectively, in the absence of IPTG (Fig. 3F). Correspondingly, pCW17 was shown to produce the lowest level of adapter from this panel (Fig. 3E). As above, we noted a discrepancy between the predicted and empirical promoter strengths: while pCW17 empirically produced the lowest levels of adapter, its promoter strength was predicted to fall in the middle of the panel based on sequences of −10 and −35 promoter elements (Table S4; see Discussion). Together, these results confirm the inverse correlation between adapter expression and the signal produced in the B3H assay.

### Forward genetic screen for improved pAdapter construct

Given that lower levels of CI-MS2^CP^ adapter led to higher B3H signal, but that the weakest constitutive promoter still expressed more CI-MS2^CP^ than the original inducible promoter (Fig. 3E), we concluded that further optimization of the pAdapter promoter was necessary. To expedite the search for a promoter giving rise to the ideal adapter concentration, we took a forward-genetic approach to identify a variant of the construct with an optimal promoter sequence. pCW17 was chosen as a parent construct for mutagenesis libraries, as it yielded the highest B3H signal for both ChiX and OxyS. From pCW17, we generated pAdapter plasmid libraries with randomized bases in the −35 region of the promoter and in the Shine-Dalgarno sequence (−35 and SD libraries, respectively). These regions were chosen for mutagenesis to vary adapter expression at both the transcriptional and translational level (Fig. 4A).

**Figure 4.**
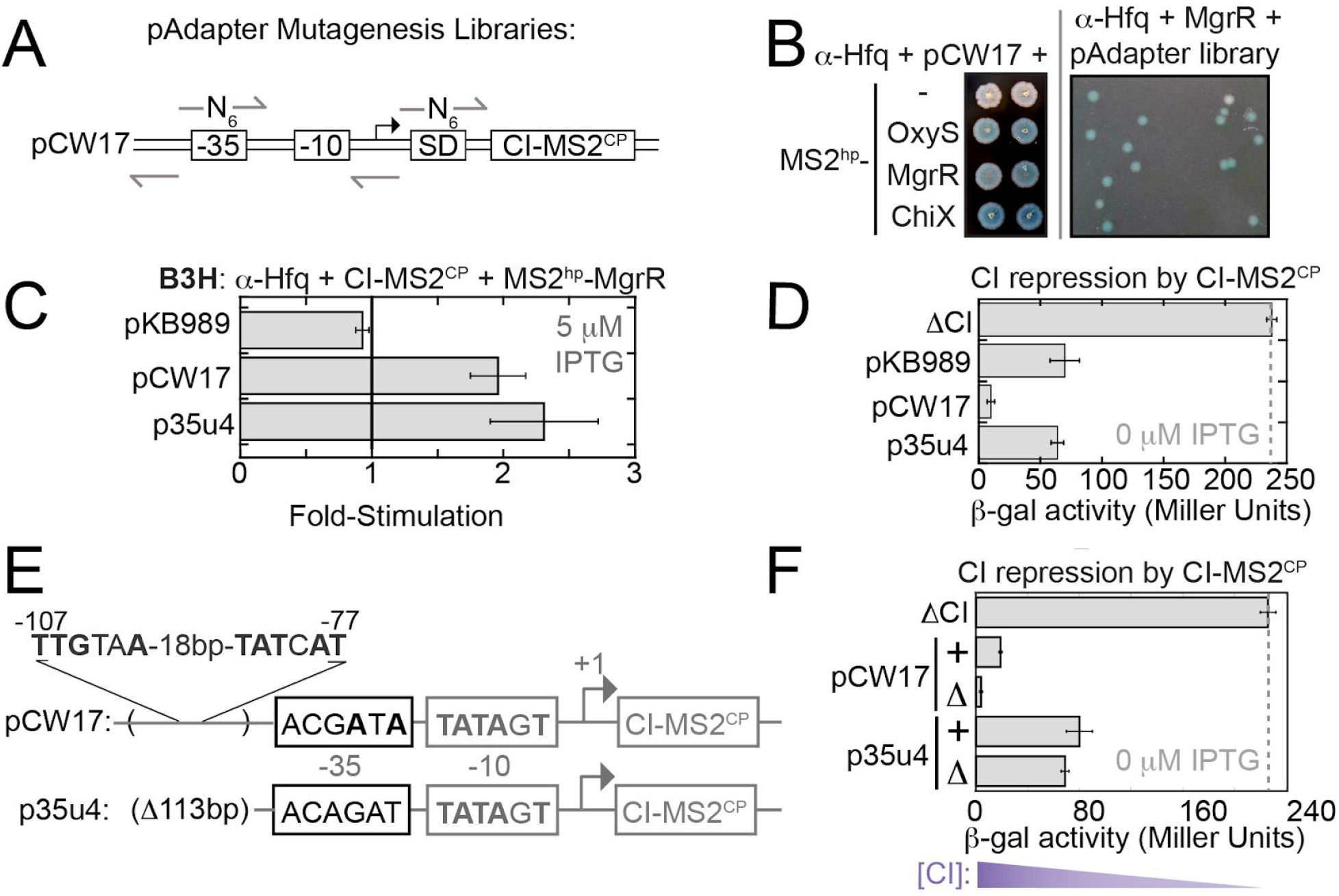
Unbiased genetic screen identifies pAdapters that produce less RNA-DNA adapter protein. (A) Schematic of mutagenesis strategy for pAdapter libraries. Six random bases (N_6_) were inserted either at the −35 promoter sequence (−35) or the core of the Shine-Dalgarno (SD) sequence in the ribosomal binding site. Arrows show the positions of DNA oligonucleotides used to construct Q5 mutagenesis (see Methods). (B) Left: Colony-color phenotypes of β-gal B3H assays between sRNAs and Hfq under conditions used for forward genetic screen. KB483 reporter cells were transformed with pAdapter (pCW17), pPrey (α-Hfq) and pBaitRNA (2xMS2^hp^-OxyS, 2xMS2^hp^-MgrR, or 2xMS2^hp^-ChiX). Right: Example plate from screen. KB473 reporter cells were transformed with pCW17-derived “-35 library”, pPrey (α-Hfq) and pBaitRNA (2xMS2^hp^-MgrR). Transformed cells were spotted on X-gal containing indicator medium (see Methods). (C) Results of B3H β-gal assays from KB483 cells grown in presence of 5 μM IPTG, with pAdapter (CI-MS2^CP^ fusion expressed from pKB989, pCW17, or p35u4), pPrey (α-Hfq) and pBaitRNA (2xMS2^hp^-MgrR). (D) Results of CI-repression β-gal assays from FW123 cells containing the indicated pAdapter plasmid. (E) DNA sequencing results from p35u4 adapter plasmid. Matches to consensus are shown in bold. The region from −54 to −166 relative to the transcription start-site (TSS) in pCW17 was missing (Δ113bp). A σ^70^ consensus match within this deleted region (from −107 to −77 relative to TSS (4/6 and 5/6 matches to consensus sequence). (F) Results of CI-repression β-gal assays from FW123 cells containing ΔCI, pCW17, pCW24 (pCW17-Δ), p35u4 or pRM24 (p35u4-Δ).

We reasoned that it would be easier to identify an optimal pAdapter sequence in the context of a relatively weak RNA-protein interaction, and chose the MgrR-Hfq interaction for this purpose. Plating conditions on X-gal containing medium were identified on which MgrR-Hfq and ChiX-Hfq interactions produced pale- and dark-blue phenotypes, respectively (Fig. 4B). To identify mutations in pAdapter that give rise to optimal CI-MS2^CP^ expression, *Δhfq* reporter cells expressing α-Hfq and MS2-MgrR were transformed with either pAdapter plasmid library (−35 or SD) and plated on X-gal containing medium to identify colonies that were more blue than the pCW17 control, indicating a stronger Hfq-MgrR interaction (Fig. 4B). Out of ~10,000 colonies screened, 40 colonies that appeared more blue than the controls were re-struck to confirm their color. Mutants that afforded bluer colonies than those with pCW17 were tested in a liquid β-gal assay; several pAdapter variants from the mutant screen indeed gave rise to fold-stimulations higher than pCW17 for Hfq-MgrR B3H interaction. One consistently promising candidate, p35u4, was found from the plasmid library with mutations introduced in the −35 region of pAdapter. This pAdapter variant produced MgrR-Hfq interactions slightly higher than pCW17 when tested in liquid assays (Fig. 4C). Based on the higher B3H signal, we expected that p35u4 would produce lower levels of adapter protein than pCW17, and confirmed this using a CI-repression assay (Fig. 4D). This indicated that the promoter in p35u4, isolated from a forward-genetic screen, is weaker than any that we generated from site-directed mutagenesis.

To identify the mutations in p35u4 that resulted in lower expression of the adapter protein, we next sequenced this plasmid. Whereas the annotated −35-promoter region of pCW17 contains 3/6 matches to the σ^70^-consensus sequence, the corresponding region of p35u4 did not possess any consensus matches. This apparently weaker −35-element is consistent with p35u4 producing less adapter than pCW17 (Fig. 4E). However, in addition to the mutations in the −35 element, the p35u4 sequence also showed an unexpected 113-bp deletion upstream of the annotated promoter (Fig. 4E). Upon closer inspection of the region of pCW17 deleted from p35u4, we identified a stronger match to the σ^70^-consensus sequence than the annotated promoter that was targeted for mutagenesis (Fig. 4E).

To test whether this upstream promoter was driving CI-MS2^CP^ expression from pCW17, we deleted the region corresponding to the downstream promoter (−35 to −6 relative to TSS) from both pCW17 and p35u4, generating “no promoter” (Δ) control plasmids. In a CI-repression assay, pCW17-Δ produced more CI-MS2^CP^ than pCW17, despite the absence of the annotated downstream promoter (Fig. 4F). In contrast, when the promoter was removed from p35u4 (Δ), adapter levels remained largely unchanged. Together, these results are consistent with read-through transcription from upstream promoters being a major source of CI-MS2^CP^ expression from these constructs (Fig. 4F; see Discussion).

### Optimal pAdapter constructs produce intermediate protein concentrations

We previously observed a roughly linear, inverse relationship between the concentration of adapter protein and the B3H signal produced for Hfq interactions with both ChiX and OxyS. To test whether this relationship remained linear across the wider range of adapter concentrations, we used a panel of 6 pAdapter candidates isolated in our forward-genetic screen, and compared the B3H signal for an MgrR-Hfq interaction to the amount of CI-MS2^CP^ generated by each pAdapter, as determined from CI-repression assays. At the highest levels of adapter expression (using pAdapters generated through site-directed mutagenesis), the relationship between MgrR-Hfq B3H signal and CI-repression is similar to previously observed (Fig. 5A). At lower adapter concentrations, however, the relationship with B3H signal begins to flatten, and more variability is observed in the relationship at the lowest levels of adapter. There appears to be an ideal concentration of adapter, after which B3H signal drops; indeed, the data are well fit by a quadratic function. p35u4, the promising candidate discussed above, is indicated in blue, and is found within the peak of the curve.

**Figure 5.**
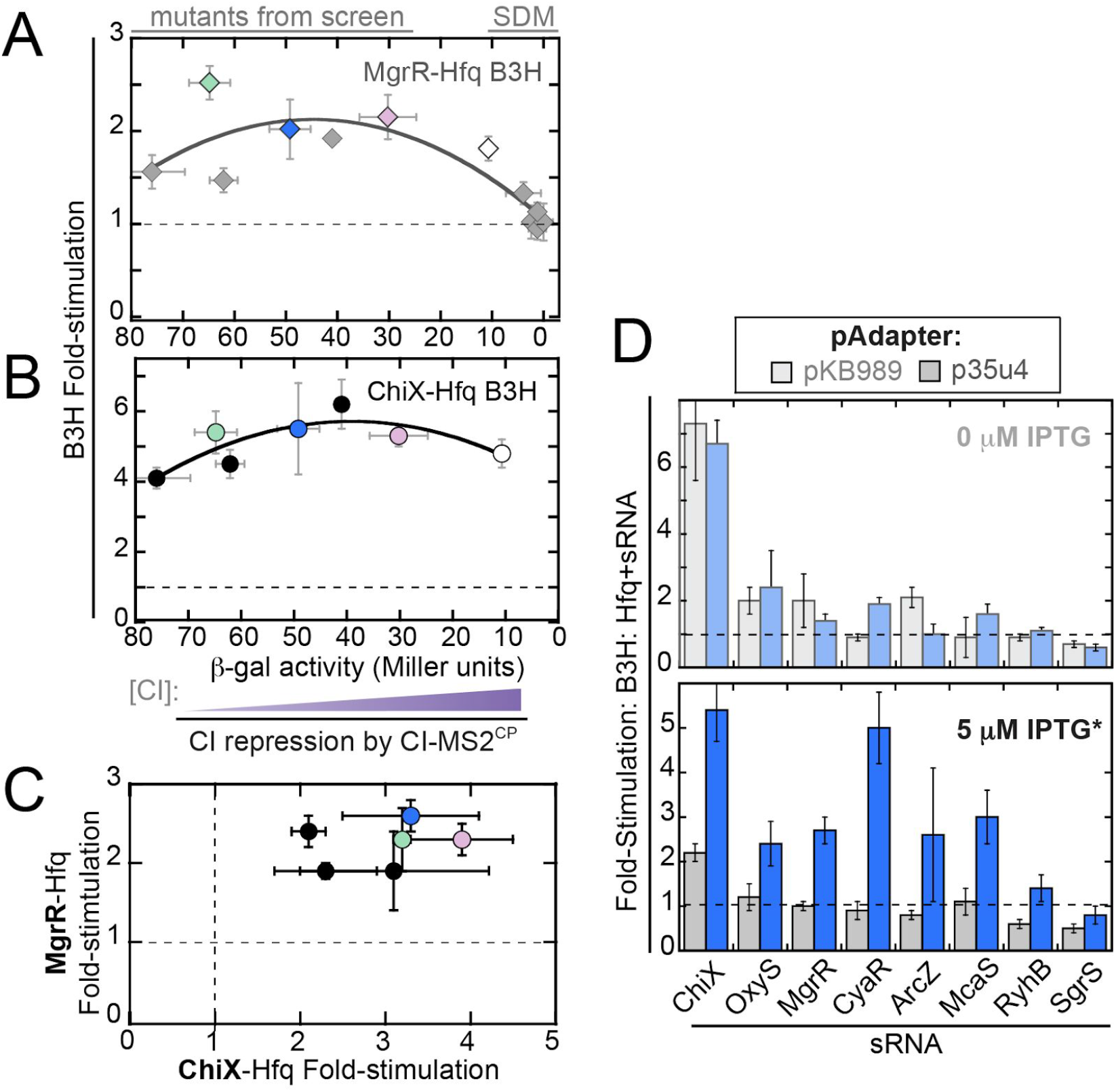
Optimal adapter expression allows for expanded detection of Hfq-sRNA interactions. (A) Effect of adapter-protein concentration on (A) MgrR-Hfq and (B) ChiX-Hfq B3H interactions. 12 pAdapter constructs are included — 6 from site-directed mutagenesis (SDM) and 6 from forward genetic screen (mutants from screen). Data points correspond to pAdapters (left to right) 35u14, p34u9 (green), SDu7, p35u4 (blue), 35u5, pSDu5 (pink), pCW17 (white), pCW18, pCW19, pCW14-16. B3H data collected with each pAdapter are plotted against CI repression data from FW123 carrying the same construct. Both data sets were collected in the absence of IPTG. Data are fit to a quadratic function (y=ax^2^+bx+c; MgrR: R=0.84; ChiX, R=0.83). (C) Comparison of pAdapter-dependence of MgrR- and ChiX-B3H interactions. Results of B3H β-gal assays performed in KB473 cells with pPrey (α-Hfq), pBaitRNA (2xMS2^hp^-ChiX or 2xMS2^hp^-MgrR) and pAdapter (left to right: pSDu7, p35u14, p35u5, p35u9 (green), p35u4 (blue), pSDu5 (pink)). (D) Results of B3H β-gal assays performed in KB473 cells with pAdapter (CI-MS2^CP^ fusion expressed from pKB989 or p35u4), pPrey (α-Hfq) and pBaitRNA (2xMS2^hp^ moieties fused to the listed sRNAs), grown in the presence of 5 μM IPTG*.

To ensure that we chose a pAdapter variant that improved B3H signal beyond the specific interaction used in the forward-genetic screen, we also measured ChiX-Hfq interactions across the same pAdapter panel. The shape of the relationship between B3H signal and CI-MS2^CP^ expression is similar between these two interactions (Fig. 5B). We next directly compared the signal for ChiX-Hfq interactions with each pAdapter to that obtained for MgrR-Hfq interactions. In this analysis, we noticed two pAdapter constructs – in addition to p35u4 – that performed well in both MgrR- and ChiX-Hfq interactions: another plasmid from the −35 library (p35u9, green) and one from the SD library (pSDu5, pink; Fig. 5C). While these plasmids produced lower levels of adapter than pCW17 (Fig. 5A,B), their sequences suggest that this occurs through distinct mechanisms from producing p35u4. An additional 118bp of DNA were present in the sequence of p35u9, relative to pCW17. This region, inserted between the upstream and downstream promoters, contains an additional strong match to the σ^70^-consensus sequence (Fig S2; see Discussion). While the −35-region of pSDu5 was unchanged relative to the parent construct, the Shine-Dalgarno region of pSDu5 contained only a one-base match to the consensus ribosomal-binding site (ACGUCAA*C*C) in contrast to the four-base match of pCW17 (AC<u>AGGA</u>AAC; Fig S2). In summary, these results show that lowering adapter expression either at a transcriptional or post-transcriptional level can increase B3H signal.

### p35u4 improves B3H signal across many Hfq-sRNA interactions

In order to test whether the increases in signal with p35u4 would be general across multiple B3H interactions, we next examined interactions between Hfq and a larger panel of sRNAs. Eight Hfq-dependent sRNAs were chosen — ChiX, McaS, OxyS, MgrR, CyaR, ArcZ, RyhB, and SgrS — which showed a range of signals in B3H interactions with Hfq from high to undetectable using pKB989 as a source of adapter (Fig 1E). In the absence of IPTG, p35u4 resulted in similar signal strength as pKB989 across this group of sRNAs (Fig 5C, top). In the presence of 5 μM IPTG, however, B3H assays conducted with p35u4 showed a substantial increase in signal for most sRNAs tested (Fig 5C, bottom). Thus, when the adapter protein is tuned to an appropriate level, induction of α-Hfq prey leads to higher B3H signal. These data demonstrate that p35u4 improves signal for many B3H interactions beyond the MgrR-Hfq interaction through which it was identified.

### Relationship between binding energetics and B3H signal

An important question is how B3H interaction signal compares to the *in vitro* binding affinity of an RNA-protein interaction. In order to test this relationship while changing as few variables as possible, we used a panel of Hfq variants, the binding affinities of which had been carefully measured *in vitro* with RNAs such as OxyS, DsrA and poly(A) (Table S5).(Mikulecky et al. 2004; Olejniczak 2011) We first established that each of these Hfq variants is stably expressed in the context of the α-Hfq fusion protein. To do so, we used the Hfq-Hfq “self” B2H interaction (Fig 1B,D).(Berry and Hochschild 2018) In a B2H assay with CI-Hfq^WT^ as bait, each of the α-Hfq variants stimulated *lacZ* transcription to a similar extent as Hfq^WT^. This suggests that each variant is stably expressed in the cell (Fig. 6A).

**Figure 6.**
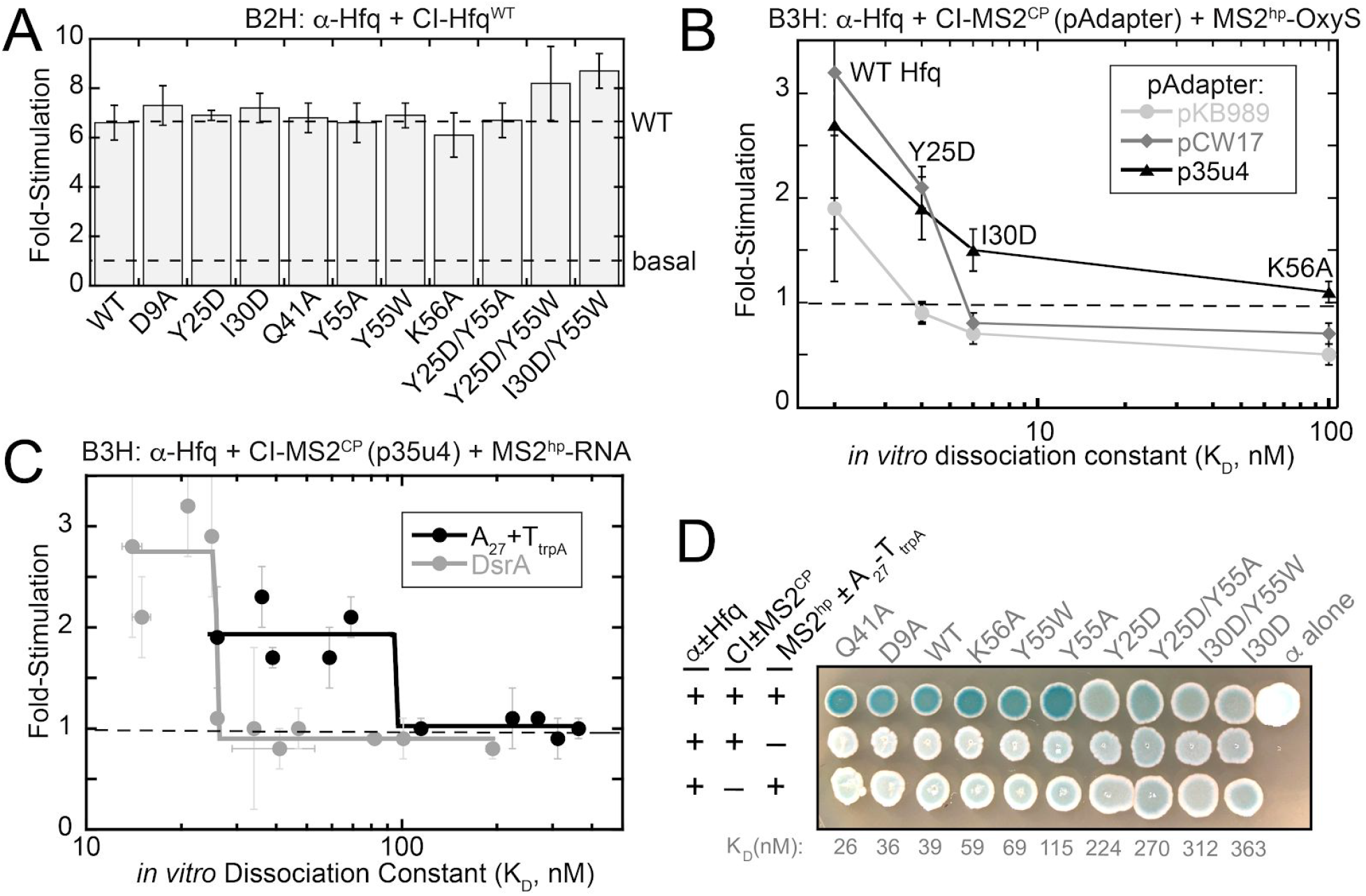
Relationship between *in vivo* B3H signal and *in vitro* binding energetics. **(A)** Results of β-gal B2H assays with KB483 reporter cells containing two compatible plasmids: one that encoded λCI or the CI-Hfq fusion protein (WT), another that encoded α or an α-Hfq fusion protein (WT or the specified mutant). (B) Results of β-gal B3H assays between OxyS and Hfq mutants. Assays were conducted in KB483 cells grown in presence of 5 μM IPTG*, with pAdapter (CI-MS2^CP^ fusion expressed from pKB989, pCW17, or p35u4), pPrey (α-Hfq; WT, Y25D, I30D, K56A) and pBaitRNA (1xMS2^hp^-OxyS). Fold-stimulation for B3H assays between OxyS RNA and each Hfq variant was plotted against the K_D_ value determined previously for each interaction using filter-binding assays.(Olejniczak 2011) (C) Results of β-gal B3H assays between A_27_ and DsrA RNAs and Hfq mutants. Assays were conducted in KB483 cells grown in presence of 5 μM IPTG*, with pAdapter (p35u4), pPrey (α-Hfq; WT or indicated variant) and pBaitRNA (2xMS2^hp^-DsrA: pHL34 or 1xMS2^hp^-A_27_-T_trpA_: pHL26). Fold-stimulation for B3H assays between both RNA and each Hfq variant was plotted against the K_D_ value determined previously for each interaction using gel-shift assays.(Mikulecky et al. 2004) (D) Colony-color phenotypes of β-gal B3H assays between A_27_ RNA and Hfq mutants. KB483 reporter cells were transformed with pAdapter (p35u4), pPrey (α-Hfq; WT or indicated variant) and pBaitRNA (1xMS2^hp^-A_27_-T_trpA_) Transformed cells were spotted on X-gal continuing indicator medium (see Methods). Negative control transformations with plasmids lacking A_27_ RNA or MS2^CP^ are shown in bottom two rows and labeled as in Fig 1C.

We first utilized a set of four Hfq variants for which published dissociation constant (K_D_) values were available for interaction with OxyS RNA. These K_D_ values ranged from 2-100 nM, when measured with filter-retention assays.(Olejniczak 2011) We compared signal for each Hfq variant when the B3H assay was conducted with the original pAdapter (IPTG-inducible, pKB989) or two constitutive pAdapter constructs (pCW17 and p35u4). The original IPTG-inducible pAdapter, pKB989, produced *lacZ* signal above background only for the strongest interaction (WT; K_D_=2nM; Fig. 6B). Assays conducted with either constitutive pAdapter construct not only resulted in higher signal for the interaction between OxyS and Hfq^WT^, but also detected the interaction between OxyS and an Hfq variant with weaker binding affinity (Y25D; K_D_ = 4 nM; Fig. 6B). p35u4, which allows for the broadest detection of sRNA-Hfq interactions (Fig. 5C) also allowed interactions defined by K_D_ values of 6 nM and 100 nM (Hfq^I30D^ and Hfq^K56A^) to be distinguished in their B3H signal. In contrast, no signal was detectable for either of these variants when assays were performed with pCW17 or pKB989. Together, these results indicate that p35u4 allows detection of — and differentiation between — RNA-protein interactions with the widest range of K_D_ values.

Using the optimized pAdapter vector (p35u4), we further explored the relationship between *in vitro* K_D_ values and *in vivo* B3H signal across a larger range of interaction strengths. We used two RNAs: the sRNA DsrA and the sequence A_27_. Binding affinities of these RNAs had been previously determined with numerous Hfq variants using gel-shift assays.(Mikulecky et al. 2004) As above, all Hfq variants performed comparably to Hfq^WT^ in a B2H assay with CI-Hfq^WT^ (Fig. 6A). We measured B3H signal using p35u4 for interactions between these RNAs and ten Hfq variants, and plotted the fold-stimulation values against the corresponding K_D_ value for each Hfq variant with either RNA (Fig. 6C; Fig S3). Overall, there is a strong but non-linear relationship between fold-stimulation and K_D_: the Hfq variants with the strongest binding affinities *in vitro* show relatively high and roughly consistent B3H signal for both A_27_-Hfq and DsrA-Hfq interactions. For each RNA, there is a K_D_ threshold at which interactions drop sharply; all interactions defined by K_D_ values beyond this threshold value remain at basal levels (Fig. 6C). For DsrA, the threshold value appears to be ~25 nM, whereas for poly(A), the threshold value appears to be around 100 nM.

In addition to β-gal assays in which activity is quantified from lysed cells that were grown in liquid culture, the B3H system can also be implemented with a plate-based assay on X-gal containing medium that provides qualitative information about RNA-protein interactions (*e.g.* Fig 4B).(Berry and Hochschild 2018; Stein et al. 2020). In order to determine whether a similar relationship between B3H signal and K_D_ would be observed in this qualitative version of the assay, we conducted plate-based assay for A_27_ interactions with the same set of Hfq variants as used in liquid assays above. Interactions defined by K_D_ values between 25 and 115 nM (Hfq^Q41A^-Hfq^Y55A^) produced bacterial patches that were similarly blue to one another, and all more blue than negative-control patches (Fig. 6D). For Hfq variants with K_D_ values above 115 nM — a similar threshold-value as observed for the liquid assays — the blueness of patches becomes comparable to the respective negative controls. Together, these results demonstrate that there is a positive relationship between the strength of an RNA-protein interaction *in vitro* and the signal obtained in our genetic B3H assay, both in quantitative liquid- and qualitative plate-based assays.

## DISCUSSION

Robust genetic systems for monitoring biomolecular interactions hold great promise for the ongoing discovery and analysis of novel RNA-binding proteins and their interacting RNAs. The bacterial three-hybrid (B3H) system relies on a network of DNA, RNA and protein molecules interacting to generate a genetic output corresponding to the strength of an RNA-protein interaction. In this work, we have systematically investigated the relationship between the intracellular concentration of one of these protein components – the RNA-DNA adapter protein (CI-MS2^CP^) – and the signal generated by the assay. We have identified an improved pAdapter construct that increases B3H signal for numerous protein-RNA interactions in this system and, importantly, extends the range of interactions that can be successfully detected by this assay into those defined by dissociation constants in the tens-of-nanomolar range. By improving assay signal and defining more clearly its relationship to *in vitro* binding affinity, we anticipate this tool will be more broadly useful to others in the bacterial RNA community.

### Successful optimization of pAdapter construct

At the outset of this work, only ~50% of well-established Hfq-sRNA interactions were detectable in our B3H system (Fig. 1E). Our strategy to perform an *in vivo* titration of the adapter protein evolved from the observation that, somewhat counterintuitively, higher levels of IPTG decreased signal in the B3H assay. The major innovation from this work is the development of p35u4 as an optimized pAdapter construct. Using this plasmid as a constitutive source of the CI-MS2^CP^, the majority of Hfq-sRNA interactions can now be detected in the B3H system (Fig. 5C). In considering why constitutive expression of the adapter from p35u4 improves signal over the original IPTG-dependent pAdapter construct, we have generated the model depicted in Fig. S4. In this model, low IPTG results in low signal with either the original or optimized pAdapter, due to low expression of prey protein. The addition of IPTG is necessary to induce α-prey fusion protein so that sufficient RNAP molecules have prey protein attached. However, in the original system, induction of adapter occurs concomitantly with induction of prey; their expression cannot be separated, and high levels of adapter may inhibit B3H signal. The introduction of a constitutive pAdapter in the optimized system allows for prey protein to be induced while keeping adapter levels constitutively low. This means that α-prey fusion protein can be titrated independently of the adapter to find the ideal level for any given prey protein. We anticipate this will allow for greater generalizability of the assay, especially as toxicity of α-prey expression can vary considerably from prey to prey.

One important question is how generalizable the improvements described here will be to other RNA-protein interactions. Conducting B3H assays with p35u4 certainly improved detection for Hfq-sRNA interactions: the optimized system allowed six out of eight sRNA-Hfq interactions to be reliably detected, when previously only four could be. In addition, the fact that the relationship between B3H signal and adapter concentration was similar in shape between MgrR- and ChiX-Hfq interactions suggests that the optimized level of adapter expression will be generalizable to multiple RNA-protein interactions. The plateau of B3H signal in relation to the concentration of CI-MS2^CP^ adapter likely represents signal becoming limited by another variable in the system. It will be important to investigate the dependence of signal on other assay parameters in the future. One lesson that can be learned from the current study is that a forward genetic screen can be very helpful to broadly explore parameter space.

### Considerations for Hybrid RNAs

For our mutant analysis, we established Hfq-RNA B3H interactions for RNAs that had been carefully studied with *in vitro* binding assays with a variety of Hfq site-directed variants. For these purposes, we used the RNAs OxyS, A_27_ and DsrA.(Mikulecky et al. 2004; Olejniczak 2011) We had previously detected OxyS-Hfq assays in the B3H assay,(Berry and Hochschild 2018) but since some effort was required to identify successful hybrid RNA sequences for the other RNAs, we explain our considerations here.

Poly(A) sequences have been used extensively in the literature to model distal face interactions with Hfq.(Sun and Wartell 2006; Link et al. 2009) In our studies with poly(A) in the B3H assay, we were unable to produce the exact A_27_ RNA used in biochemical studies,(Mikulecky et al. 2004; Olejniczak 2011) since the hybrid RNA is generated inside the cell using the cellular RNAP. The available options were to provide an intrinsic terminator at the end of the A_27_ sequence or to allow for eventual termination from a rho-dependent terminator. We chose the former strategy because it would produce a consistent and discrete RNA species. We appended the terminator from the *trpA* transcript (T_*trpA*_) as it is one of the shortest naturally occurring intrinsic terminators, and found that an A_27_ construct with T_*trpA*_ at its 3’ end gave more consistent results than a hybrid-RNA construct lacking T_*trpA*_. One important question is whether the addition of the 28-nt T_*trpA*_ sequence would influence the interactions of the A_27_ RNA with Hfq in our assay, therefore skewing the relationship between B3H signal and *in vitro* binding affinity measured between A_27_ alone. Indeed, we wondered whether the addition of T_*trpA*_ could introduce a non-native proximal-face interaction with Hfq. Two lines of evidence suggest that a proximal-face interaction with T_*trpA*_ is not a major driver of the B3H interaction between Hfq and A_27_-T_*trpA*_. First, when tested alone, the T_*trpA*_ sequence itself does not interact with Hfq in a B3H assay,(Pandey et al. 2020) though it could potentially interact in the context of the longer A_27_-containing construct. Furthermore, intrinsic terminators typically interact with Hfq through their U_6_ sequence and the proximal face. The fact that signal for interaction between A_27_ -T_*trpA*_ and Hfq is not impacted by proximal face substitutions (*e.g.* K56A, Y55W; Fig. S3B) also indicates that the *trpA* terminator is not a primary contributor to the detected interaction.

The construct used for DsrA differed from other hybrid RNAs used here in that the XmaI restriction site used to introduce the sRNA was removed from the final hybrid RNA construct. This choice was made because secondary-structure predictions suggested that the GC-rich XmaI sequence would cause the original hybrid RNA to misfold. Without this modification to the bait RNA, we were unable to reliably detect the DsrA-Hfq interaction, even with the optimized pAdapter construct p35u4. This modification of the DsrA bait RNA suggests a path forward for optimization of other sRNAs in the assay.

### Dependence of B3H signal on binding energetics

Importantly, in this work we have been able, for the first time, to quantitatively explore the relationship between B3H signal and *in vitro* binding energetics. Not only does p35u4 increase B3H signal for many Hfq-sRNA interactions, but it does so in a way that is consistent with binding affinity. This allows for differentiation between interactions with a broader range of K_D_ values than was previously possible. The shape of this relationship for interactions of both A_27_ and DsrA with Hfq demonstrates a threshold-behavior, in which signal is “high” for interactions with K_D_ values below a certain threshold value, and returns to “basal” levels beyond this threshold K_D_ value (Fig. 6C). Since DsrA and poly(A) bind to different surfaces of Hfq (Fig. S3), the identity of the Hfq variants in the “strong-binding regime” for each RNA are different; this demonstrates that it is binding affinity rather than some other feature intrinsic to each Hfq variant that corresponds to its signal in the B3H assay. We envision the threshold-K_D_-value as representing a certain weakness of interaction at which other limiting factors overwhelm the ability of the system to activate *lacZ* transcription. As the assay is further optimized and other limiting factors are diminished, the threshold-value might shift towards higher K_D_-values, allowing detection of weaker interactions.

It is interesting to note that the threshold-behavior is less apparent when Hfq-OxyS interactions are assessed with our optimal pAdapter. Instead, the data for this sRNA demonstrate a more incremental decrease in B3H signal as the *in vitro* binding affinity of an Hfq variant weakens (Fig. 6B). The difference in the shape of this curve in contrast to the threshold behavior of Hfq’s interactions with A_27_ and DsrA could be related to the fact that K_D_ values were determined with different *in vitro* methodologies (gel-shift assays for affinities with A_27_ and DsrA and filter-binding assays for affinities with OxyS).(Mikulecky et al. 2004; Olejniczak 2011) It is also striking that the threshold K_D_ value is different for A_27_-Hfq interactions (69-115 nM) than for DsrA-Hfq interactions (25-26 nM). This difference indicates that, while binding affinity of an RNA-protein interaction is one factor that contributes to B3H signal, it is not the sole determinant of signal in this system. Other contributing variables could include *(i)* geometrical constraints (*e.g.* interactions with the distal-vs. proximal faces of Hfq), *(ii)* differential turnover of RNAs inside cells, or *(iii)* folding of RNAs with MS2^hp^ and/or T. It will be interesting to further explore these variables in the future.

### Comparison to yeast genetic system

The relationship between *in vitro* binding affinity and genetic signal in the yeast-three hybrid (Y3H) assay has also been carefully assessed, and it is interesting to compare the signal and sensitivity of the two systems. The signal obtained in the B3H assay, even with the optimized system, is substantially lower than that reported for Y3H assays.(Koh and Wickens 2014a, 2014b) β-gal activity in the Y3H system spanned two orders of magnitude for interactions spanning a range of K_D_ values between 10 and 80 nM.(Hook et al. 2005) Across this range of dissociation constants, *lacZ* signal was found to be directly related to RNA-protein binding energetics. In our B3H assay, we tested RNA-protein interactions over a larger range of K_D_ values (20-350 nM) and observed β-gal activities that varied only ~4-fold. Despite the lower signal-to-noise of the B3H assay, when A_27_-Hfq data are plotted in the same manner as previous Y3H data (Fig. S5A), we observe a similarly linear relationship between the K_D_ value measured *in vitro* for a given RNA-protein interaction and the logarithm of detected β-gal activity (Fig S5B).

In previous studies with the Y3H system, signal was found to depend on the length of hybrid RNA: shorter RNAs yielded stronger interactions than longer RNAs and 150-200 nts was the optimal length of hybrid RNAs.(Bernstein et al. 2002) We note that, of all the Hfq-interacting sRNAs we have tested our this system, SgrS is the longest (227 nt). The length of this hybrid may impede signal, as was seen in the Y3H system, as SgrS is one of two sRNAs (along with RyhB) that produce very modest signal in Hfq interactions even in the optimized B3H system presented here. The length of SgrS may make it more likely to misfold and/or be degraded inside the cell.

### Transcriptional and translational control of adapter concentration

In our initial attempts to generate a new pAdapter with site-directed mutagenesis, we modulated transcription of the CI-MS2^CP^ gene by mutating its promoter region. Once we decided to take an unbiased forward-genetic approach, we designed libraries that would allow us to discover optimized pAdapter constructs which tuned adapter concentration through either translational or transcriptional control. Indeed, results of our forward-genetic screen demonstrated that there were multiple successful paths to tune to levels of the adapter protein. The expression of CI-MS2^CP^ could be reduced either by mutation of the ribosome-binding site (RBS; pSDu5) while leaving transcriptional control apparently unchanged or by altering the promoter region and leaving the RBS intact (p35u4 and p35u9).

Interestingly, successful mutants isolated from the −35 library were not simple substitutions in the −35-promoter element as we expected. Instead, we found plasmids that had large insertions or deletions upstream of the annotated promoter. This suggests that transcriptional regulation of CI-hybrid mRNA from the pAC-λCI vector may be more complex than previously understood. Indeed, one unanticipated outcome of this study was the identification of a previously unannotated promoter in this vector — used in B2H and B3H assays — upstream of the annotated *lacUV5* promoter (Fig. 4E). The presence of this upstream promoter may increase leaky expression of CI-fusion proteins in B2H and related assays. Indeed, we envision that, in our first round of constitutive pAdapter constructs, weak downstream promoters may have acted as inhibitory elements, perhaps blocking transcriptional read-through from the stronger upstream promoter. This model helps to rationalize why apparently weaker promoter sequences in our site-directed mutagenesis efforts resulted counterintuitively in stronger expression of the adapter protein (*e.g.,* Fig. 3E).

### Outlook

We have previously used the B3H assay in both forward- and reverse-genetic setups to probe molecular mechanisms of ProQ- and Hfq-RNA interactions.(Berry and Hochschild 2018; Pandey et al. 2020) Here, we have utilized a similar strategy of forward- and reverse-genetic approaches, but have applied these tools to improving the operation of the assay itself.

In the course of this study, the signal-to-noise in the B3H system has been improved for interactions between several sRNAs and the Hfq protein. Despite promising advances, there is still certainly room for improvement in B3H signal. Even with the optimizations made here, the assay only reliably detects interactions defined by dissociation constants 10 nM or stronger, with no demonstrated successes for interactions with K_D_ values higher than 80 nM. While there are many high-affinity RNA-protein interactions that fall within this low-nanomolar range, there are also many physiologically relevant RNA-protein interactions with weaker binding affinities.(*e.g.* Yang et al. 2013)

In order to further increase the robustness of the assay, we will continue to systematically explore its dependence on multiple features, including the sequence of the hybrid RNA and the position of the O_L_2 site relative to the test promoter. Given the non-intuitive effects of cellular concentrations on signal highlighted by this study, computational modeling may also be a useful path for future optimization. In addition to providing a more robust pAdapter construct for future B3H studies, we anticipate that the findings shared here provide a helpful roadmap for future optimizations through a systematic exploration of parameter space. If genetic systems present complexities as hurdles to overcome, they also provide facile paths to overcome them through genetic screens.

## SUPPLEMENTAL MATERIAL

Supplemental material is available for this article.

## ACKNOWLEDGEMENTS

We thank Amy Camp, and members of the Berry, Camp and Lijek laboratories for advice and discussion. This work was supported by the National Institutes of Health [R15GM135878 to KEB]; and the Henry R. Luce foundation; and Mount Holyoke College.

