## Supplemental Materials for "Optimization of a bacterial three-hybrid assay through *in vivo* titration of an RNA-DNA adapter-protein"

This pdf file includes:

Supplemental Figures S1 to S5

Supplemental Tables S1 to S5

Supplemental References

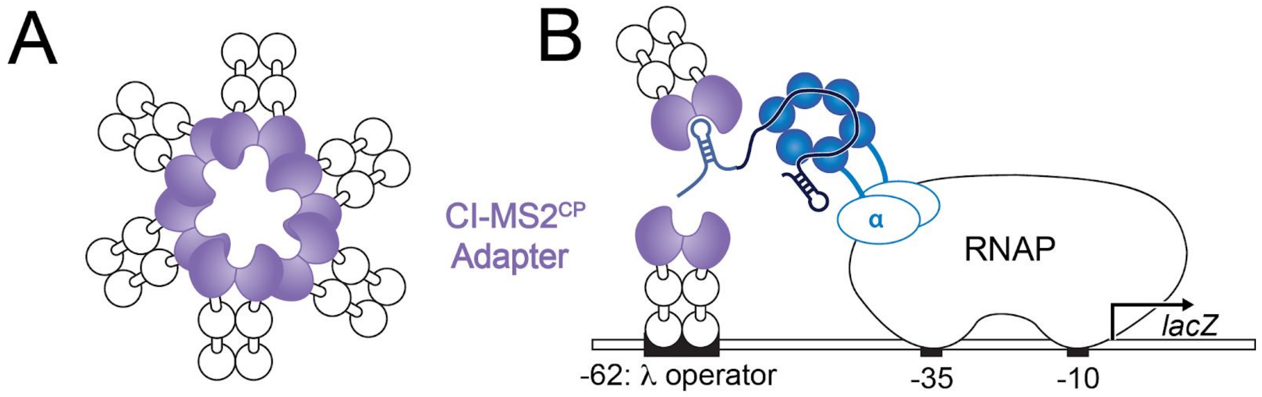


**Supplemental Figure 1.** Possible models for why B3H signal would be inhibited by high concentrations of adapter protein (CI-MS2$^{CP}$). (A) While we are using a multimerization-resistant version of MS2$^{CP}$, (Macías et al. 2008) this component is still derived from a viral coat protein that has the potential to form large, higher-order assemblies in a concentration-dependent manner. It is possible that the adapter protein could multimerize at high concentrations through its MS2$^{CP}$ moiety, in a way that blocks effective assembly of the B3H components. (B) Even in the absence of adapter multimzeriation, high concentrations of the adapter could inhibit B3H signal by "capping" hybrid RNAs. That is, if the adapter is present at high enough concentrations for one copy to bind the test promoter and a separate copy to bind the MS2$^{hp}$ moiety of the hybrid RNA. The adapter would no longer serve as an effective bridge to stabilize RNAP at the test promoter.

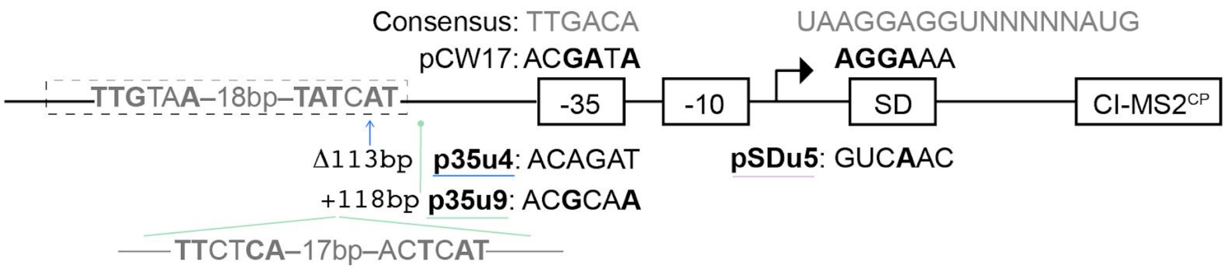


**Supplemental Figure 2.** Schematic depicting sequences of improved pAdapter constructs identified in forward genetic screen. The consensus sequence for *E. coli* $\sigma^{70}$ promoter element (-35) along with the Shine-Dalgarno (SD) sequence are shown along the top in grey. The sequences for each of these elements in pCW17 are indicated above the line schematic. Matches to consensus are indicated with bold lettering. Below the line schematic are shown the sequences of three pAdapter constructs isolated from the pCW17-derived mutagenesis library (p35u4, p35u9 isolated from a plasmid library in which the -35 region was mutated and pSDu5 from one in which the Shine Dalgarno element was mutated). Regions of these plasmids in which the sequence differs from the parental pCW17 sequence are shown. Matches to consensus are indicated with bold lettering. pSDu5 only differed from pCW17 in the SD region, whereas p35u4 and p35u9 differed from pCW17 in the -35 region, as well as in an upstream deletion (Δ113bp; p35u4) or insertion (+118bp; p35u9) relative to pCW17. $\sigma^{70}$ promoter matches in the deleted region (dashed box) or in the inserted region are shown.

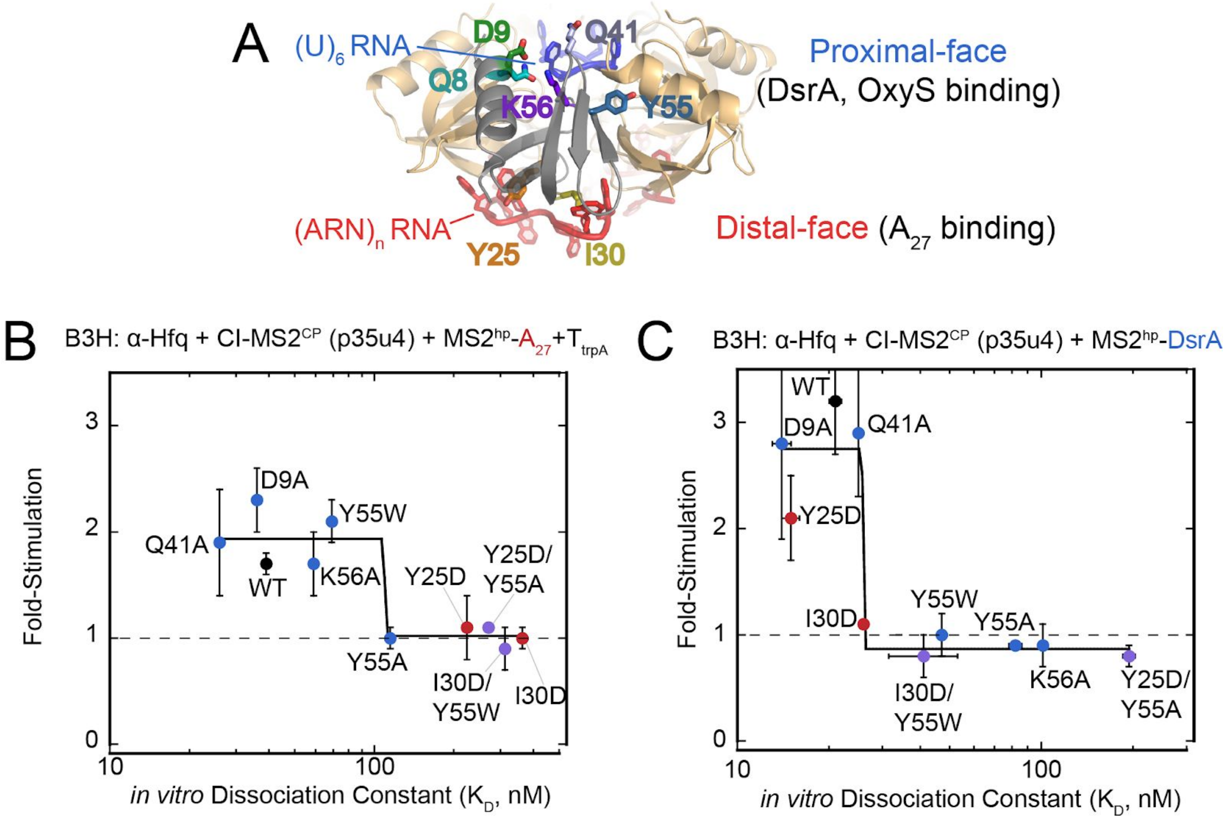


**Supplemental Figure 3.** Location of Hfq residues targeted for mutagenesis. (A) Crystal structure of Hfq hexamer, bound to poly(U) and poly(A) RNA on the proximal and distal faces, respectively (PDB ID = 4HT9).(Wang et al. 2013) One hexamer is shown as a grey cartoon; the side chains of residues targeted for mutagenesis are shown on this grey monomer in stick representation. (B,C) Results of β-gal B3H assays between $A_{27}$ and DsrA RNAs and Hfq mutants. Assays were conducted in KB483 cells grown in presence of 5 uM IPTG*, with pAdapter (p35u4), pPrey (α-Hfq; WT or indicated variant) and pBaitRNA (2xMS2$^{hp}$-DsrA, pHL34 or 1xMS2$^{hp}$-$A_{27}$-$T_{trpA}$, pHL26). Fold-stimulation for B3H assays between both RNA and each Hfq variant was plotted against the $K_D$ value determined previously for each interaction using gel-shift assays. (Mikulecky et al. 2004) Data for (B) $A_{27}$ and (C) DsrA from Figure 6C are plotted on separate axes with each individual Hfq mutant labeled. Data points are colored based on the surface of Hfq that is affected by the mutation (red = distal; blue = proximal; purple = double mutant with substitutions on both faces).

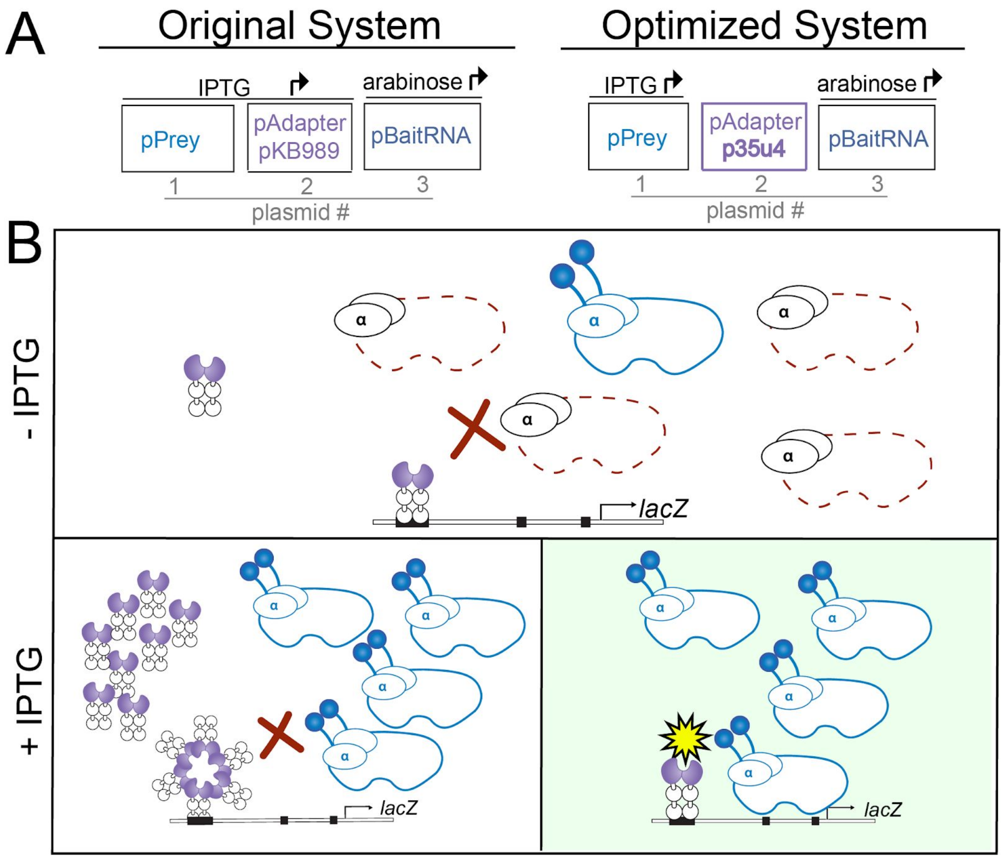


**Supplemental Figure 4.** Model for how a constitutive pAdapter, such as p35u4, improves B3H assay signal. (A) Schematic showing the plasmids used to express prey, adapter, and bait hybrid components in the original vs. optimized B3H system, along with inducers that drive expression of each component. The key difference is that adapter protein was originally induced alongside prey fusion protein with IPTG; in the optimized system, adapter concentration is fixed through a constitutive pAdapter construct, and the concentration of prey fusion protein can be modulated independently. (B) Depiction of relative concentrations of assay components in the absence (top) or presence (bottom) of IPTG when assay is conducted with original pAdapter (IPTG-inducible, pKB989, left) or with optimized pAdapter (constitutive, p35u4, right). In the absence of IPTG, both prey and adapter proteins are expressed at low levels (through leaky and/or constitutive expression). Signal is limited by the low concentration of prey fusion protein, such that most RNAP molecules in the cell are not displaying the prey fusion protein (shown as dashed red outlines). In the presence of IPTG, more prey fusion protein is expressed, leading to more RNAP molecules displaying prey (shown in blue). However, high concentrations of the adapter are also achieved in the original system, which may inhibit B3H signal (see Fig S1). In the optimized system and in the presence of IPTG (bottom, right), adapter expression is kept low while prey expression is high, leading to optimal signal.

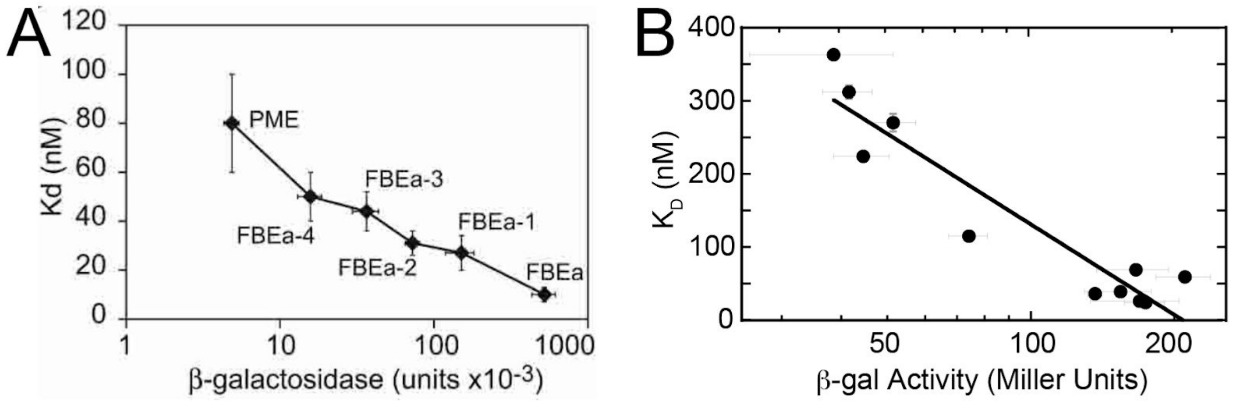


**Supplemental Figure 5.** Comparison of relationship between *in vitro* binding affinities and genetic signal in yeast three-hybrid (Y3H) and B3H systems. (A) Panel is reproduced from Hook et al. 2005. Comparison of *in vitro* binding affinities measured by gel-shift assays to Y3H reporter gene activation in YBZ1 for interactions between the PUF protein FBF-1 and six RNAs.(Hook et al. 2005) (B) Data from Figure 6C plotted in the same manner as Y3H data in (A). Results of β-gal B3H assays between $A_{27}$ RNA and Hfq mutants conducted in KB483 cells grown in presence of 5 uM IPTG* with pAdapter (p35u4), pPrey (α-Hfq; WT or indicated variant) and pBaitRNA (1xMS2$^{hp}$-A27-$T_{trpA}$, pHL26). The $K_D$ value for interaction between $A_{27}$ RNA and each Hfq variant — determined previously using gel-shift assays (Mikulecky et al. 2004) — was plotted against raw β-gal value activity measured for each interaction, plotted on a logarithmic scale. Data are fit to the logarithmic function y=a+b*log(x), R=0.94.

**SUPPLEMENTAL TABLES**

**Table S1.** *E. coli* strains used in this study.

| Strain | Genotype | Antibiotic Resistance | Source | Figures |
|---|---|---|---|---|
| NEB 5α- F'$I^q$ Competent *E. coli* | *E. coli* host strain for plasmid construction: F´ *proA*$^+$*B*$^+$ *lacI*$^q$ Δ*(lacZ)M15 zzf::Tn10* (Tet$^R$) / *fhuA2*Δ*(argF-lacZ)U169 phoA glnV44 Φ80Δ(lacZ)M15 gyrA96 recA1 relA1 endA1 thi-1 hsdR17* | TetR | New England Biolabs | [used for cloning] |
| KB473 | FW102 Δ*hfq*::FRT harboring F' episome bearing kanamycin resistance and test promoter *placO*$_L$*2-62* fused to *lacZ* | KanR; StrR | (Berry and Hochschild 2018) | Fig 1, 2B, Fig 3, Fig 5 |
| KB483 | FW102 *hfq::kan* harboring F' episome bearing tetracycline resistance and test promoter *placO*$_L$*2-62* fused to *lacZ* | TetR; StrR | (Pandey et al. 2020) | Fig 2C, Fig 6 |
| FW123 | FW102 harboring F' episome bearing kanamycin resistance and a *lacZ* reporter gene controlled by a promoter harboring the λCI operator positioned between the - 35 and -10 elements. | KanR | (Pinkett et al. 2006; Whipple et al. 1994) | Fig 3B,C,E,F; Fig 4D,F; Fig 5A,B |

**Table S2. Plasmids used in this study**

| Name | Description | Details | Antibiotic Resistance | Reference/ Source | Figures |
|---|---|---|---|---|---|
| pACλCI | pACλCI-empty vector | Encodes full length λCI under control of *lacUV5* promoter | CmR | (Berry and Hochschild 2018) | through out |
| pACλCI-β-flap | pACλCI-β-flap$^{831-1057}$ | $P_{lacUV5}$ -directed synthesis of the λCI protein fused via three alanines to residues 831-1057 of the β subunit of *E. coli* RNAP | CmR | (Deighan et al. 2008) | Fig 2C |
| pACΔCI | pAC-empty vector | Comparable to pACλCI but lacking λCI ORF following *lacUV5* promoter | CmR | (Pinkett et al. 2006; Whipple et al. 1994) | Fig 3C,E |
| pBRα | pBr-α-empty vector | Encodes residues 1-248 of the alpha vector fused under control of *lpp* and *lacUV5* promoters | AmpR | (Berry and Hochschild 2018) | through out |
| pBR-σ$^{70}$ | pBR-σ70$^{D581G}$ | $P_{lacUV5}$ /$P_{lpp}$ -directed synthesis of the λCI protein fused directly to *E. coli* σ$^{70}$ region 4 (residues 528-613 of σ$^{70}$). The σ$^{70}$ moiety also carries the D581G substitution | AmpR | (Kuznedelov et al. 2002) | Fig 2C |
| pCH9 | pCDF-pBAD-1xMS2$^{hp}$-OxyS | *E. coli oxyS* inserted into pCH1 following 1xMS2$^{hp}$ between XmaI and HindIII sites; intrinsic terminator encoded by sRNA | SpecR | (Pandey et al. 2020) | Fig 6B |
| pCW14 | pAC-$p_{constit}$-CI-MS2 | Encodes CI-MS2$^{CP}$ fusion protein under the control of a constitutive promoter. Comparable to pCW18 (Δ*lacO*) but with -35: ACGATA and -10 TATACT | CmR | This study (oCW28 + oCW29 + oCW30 + oCW31) | Fig 3E,F; 5A,B |
| pCW15 | pAC-$p_{constit}$-CI-MS2 | Encodes CI-MS2$^{CP}$ fusion protein under the control of a constitutive promoter. Comparable to pCW18 (Δ*lacO*) but with -35: CCGATA and -10 TATACT | CmR | This study (oCW30 + oCW31) | Fig 3E,F; 5A,B |
| pCW16 | pAC-$p_{constit}$-CI-MS2 | Encodes CI-MS2$^{CP}$ fusion protein under the control of a constitutive promoter. Comparable to pCW18 (Δ*lacO*) but with -35: CCGACA and -10 TATACT | CmR | This study (oCW32 + oCW33+ oCW30 + oCW31) | Fig 3E,F; 5A,B |
| pCW17 | pAC-$p_{constit}$-CI-MS2 | Encodes CI-MS2$^{CP}$ fusion protein under the control of a constitutive promoter. Comparable to pCW18 (Δ*lacO*) but with -35: ACGATA and -10 TATAGT | CmR | This study (oCW28 + oCW29) | Fig 3E,F; 4B, C, D, F; 5A,B; 6B |

| | | | | | |
|---|---|---|---|---|---|
| pCW18 | pAC-$p_{constit}$-CI-MS2 | Encodes CI-MS2$^{CP}$ fusion protein under the control of a constitutive promoter. Relative to *lacUV5* promoter in pKB989, Δ*lacO*, -35: CCGACA, -10 TATAGT | CmR | This study; oCW32 + oCW33) | Fig 3C,D,E, F; 5A,B |
| pCW19 | pAC-$p_{constit}$-CI-MS2 | Encodes CI-MS2$^{CP}$ fusion protein under the control of a constitutive promoter. Comparable to pCW18 (Δ*lacO*) but with -35: CCGATA and -10 GATAGT | CmR | This study (oCW34 + oCW35 | Fig 3E,F; 5A,B |
| pCW24 | pAC-$p_{constit}$-CI-MS2-Δpromoter | Comparable to pCW17, encodes the CI-MS2$^{CP}$ fusion protein, but 30bp which encoded the constitutive promoter (-10 to -35 elements) have been deleted. | CmR | This study (oCW45 + oCW46 on pCW17) | Fig 4F |
| pHL6 | pCDF-pBAD-1xMS2$^{hp}$-$T_{trpA}$ | Intrinsic terminator from *trpA* operon cloned behind MS2$^{hp}$ and HindIII site in pCH1 | SpecR | (Pandey et al. 2020) | Fig 6C, D; S3; S5 |
| pHL23 | pBr-α-Ec-Hfq-I30D | *hfq* I30D mutation introduced to pKB817 by PCR | AmpR | This study (oHL52 + oHL53) | Fig 6A-D |
| pHL26 | pCDF-pBAD-1xMS2$^{hp}$-$A_{27}$-$T_{trpA}$ | poly(A) sequence of 27 adenosine nucleotides cloned behind MS2$^{hp}$ between XmaI/HindIII residues in pHL6; transcriptional termination provided by *trpA* terminator | SpecR | This study (oHL54 + oHL55 into pHL6) | Fig 6C, D; S3; S5 |
| pHL34 | pCDF-pBAD-2xMS2$^{hp}$-ΔXmaI-DsrA | *E. coli dsrA* inserted into pKB845 following 2xMS2$^{hp}$ between XmaI and HindIII sites; XmaI site removed by PCR; intrinsic terminator encoded by sRNA | SpecR | This study (oHL62 + oHL63 on pKB941) | Fig 6C; S3C |
| pHL54 | pBr-α-Ec-Hfq-D9A | *hfq* D9A mutation introduced to pKB817 by PCR | AmpR | This study (oHL88 + oHL89) | Fig 6A,C,D; S3 |
| pHL58 | pBr-α-Ec-Hfq-Q41A | *hfq* Q41A mutation introduced to pKB817 by PCR | AmpR | This study (oHL9 6+ oHL97) | Fig 6A,C,D; S3 |
| pHL60 | pBr-α-Ec-Hfq-Y55W | *hfq* Y55W mutation introduced to pKB817 by PCR | AmpR | This study (oHL100 + oHL101) | Fig 6A,C,D; S3 |
| pHL61 | pBr-α-Ec-Hfq-Y25D-Y55A | *hfq* Y25D and Y55A mutations introduced to pKB817 by PCR | AmpR | This study (oHL102 + oHL103) | Fig 6A,C,D; S3 |
| pHL63 | pBr-α-Ec-Hfq-I30D-Y55W | *hfq* I30D and Y55W mutations introduced to pKB817 by PCR | AmpR | This study (oHL100 + oHL101) | Fig 6A,C,D; S3 |

| | | | | | |
|---|---|---|---|---|---|
| pKB815 | pBr-α-MS2$^{CP}$ | Encodes residues 1-248 of alpha fused via three alanine residues to MS2 coat protein with following mutations to avoid multimerization: V30I, A81G, positions 68-80 deleted (VATQTVGGVELPV). Analogous to pKB994 (Berry and Hochschild 2018), but with a shorter linker. | AmpR | This study (oKB1038 + oKB1039) | Fig 2C |
| pKB816 | pACλCI-Hfq | Encodes residues 1-236 of λCI fused via three alanine residues to full-length wild-type *E.coli* Hfq | CmR | (Berry and Hochschild 2018) | Fig 1D; 6A |
| pKB817 | pBr-α-Hfq | Encodes residues 1-248 of alpha vector linked to full-length wild-type *E. coli* Hfq via three alanine residues; inserted into pBRα between NotI and BamHI sites | AmpR | (Berry and Hochschild 2018) | Fig 1C-E; 2B; 3D,F; 4B,C; 5A-D; 6A-D; S3; S5 |
| pKB845 | pCDF-pBAD-2xMS2$^{hp}$ | Two MS2 hairpins (2xMS2$^{hp}$) and an XmaI site inserted into pKB822 CDF origin vector between BamHI and HindIII sites | SpecR | (Berry and Hochschild 2018) | through out |
| pKB854 | pCDF-pBAD-2xMS2$^{hp}$-SgrS | *E. coli sgrS* inserted into pKB845 following 2xMS2$^{hp}$ between XmaI and HindIII sites; intrinsic terminator encoded by sRNA | SpecR | This study (oKB1107 + oKB1108) | Fig 1E; 5D |
| pKB856 | pCDF-pBAD-2xMS2$^{hp}$-MgrR | *E. coli mgrR* inserted into pKB845 following 2xMS2$^{hp}$ between XmaI and HindIII sites; intrinsic terminator encoded by sRNA | SpecR | (Berry and Hochschild 2018) | Fig 1E; 4B,C; 5A,C,D |
| pKB872 | pBr-α-Ec-Hfq-Y55A | *hfq* Y55A mutation introduced to pKB817 by PCR | AmpR | This study (oKB1117 + oKB1118) | Fig 6A,C,D; S3 |
| pKB873 | pBr-α-Ec-Hfq-K56A | *hfq* K56A mutation introduced to pKB817 by PCR | AmpR | This study (oKB1119 + oKB1120) | Fig 6A-D; S3 |
| pKB905 | pBr-α-Ec-Hfq-Y25D | *hfq* Y25D mutation introduced to pKB817 by PCR | AmpR | This study (oKB1186 + oKB1187) | Fig 6A-D; S3 |
| pKB909 | pCDF-pBAD-2xMS2$^{hp}$-ChiX | *E. coli chiX* inserted into pKB845 following 2xMS2$^{hp}$ between XmaI and HindIII sites; intrinsic terminator encoded by sRNA | SpecR | (Berry and Hochschild 2018) | Fig 1C,E; 5B-D |

| | | | | | |
|---|---|---|---|---|---|
| pKB910 | pCDF-pBAD-2xMS2$^{hp}$-McaS | *E. coli mcaS* inserted into pKB845 following 2xMS2$^{hp}$ between XmaI and HindIII sites; intrinsic terminator encoded by sRNA | SpecR | (Berry and Hochschild 2018) | Fig 1E; 5D |
| pKB911 | pCDF-pBAD-2xMS2$^{hp}$-CyaR | *E. coli cyaR* inserted into pKB845 following 2xMS2$^{hp}$ between XmaI and HindIII sites; intrinsic terminator encoded by sRNA | SpecR | This study (oKB1194 + oKB1195) | Fig 1E; 5D |
| pKB912 | pCDF-pBAD-2xMS2$^{hp}$-OxyS | *E. coli oxyS* inserted into pKB845 following 2xMS2$^{hp}$ between XmaI and HindIII sites; intrinsic terminator encoded by sRNA | SpecR | (Berry and Hochschild 2018) | Fig 1E; 5D; 6B |
| pKB913 | pCDF-pBAD-2xMS2$^{hp}$-RyhB | *E. coli ryhB* inserted into pKB845 following 2xMS2$^{hp}$ between XmaI and HindIII sites; intrinsic terminator encoded by sRNA | SpecR | (Berry and Hochschild 2018) | Fig 1E; 5D |
| pKB941 | pCDF-pBAD-2xMS2$^{hp}$-DsrA | *E. coli dsrA* inserted into pKB845 following 2xMS2$^{hp}$ between XmaI and HindIII sites; intrinsic terminator encoded by sRNA | SpecR | This study (oKB1209 + oKB1210) | n/a (cloning interme diate) |
| pKB942 | pCDF-pBAD-2xMS2$^{hp}$-ArcZ | *E. coli arcZ* inserted into pKB845 following 2xMS2$^{hp}$ between XmaI and HindIII sites; intrinsic terminator encoded by sRNA | SpecR | This study (oKB1211 + oKB1212) | Fig 1E; 5D |
| pKB989 | pACλCI-MS2$^{CP}$ | Encodes residues 1-236 of CI fused to MS2 coat protein (MS2$^{CP}$) via three alanine residues; includes mutations V30I, A81G, and deletion of residues 68-80 in order to avoid multimerization. This fusion protein is expressed under control of a *lacUV5* promoter. | CmR | (Berry and Hochschild 2018) | Fig 1B,C,E; 3C,D,E; 4C,D; 5D; 6B |
| pRM24 | pAC-p$_{constit*}$-CI-MS2-Δpromoter | Comparable to p35u4, encodes the CI-MS2$^{CP}$ fusion protein, but 30bp which encoded the constitutive promoter (-10 to -35 elements) have been deleted. | CmR | This study (oCW45 + oRM35) | Fig 4F |
| p35u4 | pAC-p$_{constit*}$-CI-MS2 | Encodes CI-MS2$^{CP}$ fusion protein under control of a constitutive promoter. Isolated from a pCW17-derived mutagenesis library. See Fig S3 for changes relative to pCW17. | CmR | This study (forward genetic screen) | 4C,D; 5A-D |
| p35u9, pSDu5 | pAC-p$_{constit}$-CI-MS2 | | | | Fig 5A-C |

**Table S3. Oligonucleotides used in the study.** QC indicates primers used for Quikchange-style mutagenesis; Q5 indicates primers designed with NEBBase changer for Q5 Site-Directed Mutagenesis.

| Name | Description | Used for | Sequence |
|---|---|---|---|
| oCW5 | F pAdapter | Sequencing primer | CAGACCAAAACGATCTCAAGAAGATCATC |
| oCW28 | F -35 ACGATA | QC PCR: pCW14, 17 | CCAGGCCTCGAGACGATAGCTAGCTCAGTC |
| oCW29 | R -35 ACGATA | QC PCR: pCW14, 17 | GACTGAGCTAGCTATCGTCTCGAGGCCTGG |
| oCW30 | F -10 TATACT | QC PCR: pCW14, 15, 16 | GCTCAGTCCTAGGTATACTGCTAGCGCATGCC |
| oCW31 | R -10 TATACT | QC PCR: pCW14, 15, 16 | GGCATGCGCTAGCAGTATACCTAGGACTGAGC |
| oCW32 | F -35 CCGACA | QC PCR: pCW16, 18 | CAGGCCTCGAGCCGACAGCTAGCTCAGT |
| oCW33 | R -35 CCGACA | QC PCR: pCW16, 18 | ACTGAGCTAGCTGTCGGCTCGAG CCTG |
| oCW34 | F -10 GATAGT | QC PCR: pCW19 | GCTCAGTCCTAGGGATAGTGCTAGCGCATGCC |
| oCW35 | R -10 GATAGT | QC PCR: pCW19 | GGCATGCGCTAGCACTATCCCTAGACTGAGC |
| oCW22 | F -35 CCGATA | QC PCR: pCW 15,19 | CAGGCCTCGAGCCGATAGCTAGCTCAGTC |
| oCW23 | R -35 CCGATA | QC PCR: pCW 15,19 | GACTGAGCTAGCTATCGGCTCGAGGCCTG |
| oCW36 | F -35 library | Q5 PCR: -35 library | AGGCCTCGAGNNNNNNGCTAGCTCAGTCCTAGG |
| oCW37 | R -35 library | Q5 PCR: -35 library | GGGGTGCCTAATGAGTGA |
| oCW40 | F SD library | Q5 PCR: SD library | CATGCCACACNNNNNNCAGCGTATGAGC |
| oCW41 | R SD library | Q5 PCR: SD library | CGCTAGCACTATACCTAG |
| oHL52 | F Hfq I30D | Q5 PCR: pHL24 | GGTGAATGGTGACAAGCTGCAAG |
| oHL54 | F $A_{27}$ | Q5 PCR: pHL26 | AAAAAAAAAAAAAAAAAAAAAGCTTAGCCCGCCTAATGAG |
| oHL55 | R $A_{27}$ | Q5 PCR: pHL26 | TTTTTTTTCCCGGGCTGCAGACATGG |
| oHL58 | R Hfq I30D | Q5 PCR: pHL24 | AAATAAATAGAAACTGGAACAC |
| oHL62 | F DsrA ΔXmaI | Q5 PCR: pHL34 | AACACATCAGATTTCCTGG |
| oHL63 | R DsrA ΔXmaI | Q5 PCR: pHL34 | CTGCAGACATGGGTGATC |
| oHL88 | F Hfq D9A | Q5 PCR: pHL54 | TCTTTACAAGCTCCGTTCCTGAACGCAC |
| oHL89 | R Hfq D9A | Q5 PCR: pHL54 | TTGCCCCTTAGCTGCGGC |
| oHL96 | F Hfq Q41A | Q5 PCR: pHL58 | GTCTTTTGATGCGTTCGTGATCCTGTTGAAAAAC |

| | | | |
|---|---|---|---|
| oHL97 | R Hfq Q41A | Q5 PCR: pHL58 | TCGATTTGCCCTTGCAGC |
| oHL100 | F Hfq Y55W | Q5 PCR: pHL60/63 | CCAGATGGTTTGGAAGCACGCGA |
| oHL101 | R Hfq Y55W | Q5 PCR: pHL60 + pHL63 | CTGACCGTGTTTTTCAAC |
| oHL102 | F Hfq Y55A | Q5 PCR: pHL61 | CCAGATGGTTGCCAAGCACGCGA |
| oHL103 | R Hfq Y55A | Q5 PCR: pHL61 | CTGACCGTGTTTTTCAACAG |
| oRM20 | R Seq pAdapter | Sequencing primer | GCTTTAAGGCGACGTGCGTCC |
| oKB1038 | F NotI MS2$^{CP}$ | PCR: pKB815 | ATAAGAATGCGGCCGCAGCTTCTAACTTTACT CAGTTCGTTCTCG |
| oKB1039 | R BamHI MS2$^{CP}$ | PCR: pKB815 | CTTCGGATCCTTAGTAGATGCCGGAG TTTGCTGC |
| oKB1117 | F Hfq Y55A | QC PCR: pKB872 | GTCAGCCAGATGGTTGCCAAGCACGCGATTTC |
| oKB1118 | R Hfq Y55A | QC PCR: pKB872 | GAAATCGCGTGCTTGGCAACCATCTGGCTGAC |
| oKB1119 | F Hfq K56A | QC PCR: pKB873 | CAGCCAGATGGTTTACGCGCACGCGATTTCTA C |
| oKB1120 | R Hfq K56A | QC PCR: pKB873 | GTAGAAATCGCGTGCGCGTAAACCATCTGGCT G |
| oKB1186 | F Hfq Y25D | QC PCR: pKB905 | CGTGTTCCAGTTTCTATTGATTTGGTGAATGGT ATTAAGC |
| oKB1187 | R Hfq Y25D | QC PCR: pKB905 | GCTTAATACCATTCACCAAATCAATAGAAACTG GAACACG |
| oKB1209 | F DsrA XmaI | PCR: pKB941 | TCCCCCCGGGAACACATCAGATTTCCTGGTGT AAC |
| oKB1210 | R DsrA HindIII | PCR: pKB941 | CCGGCCAAGCTTAAAAAAAATCCCGACCCTGA GGG |
| oKB1211 | F ArcZ XmaI | PCR: pKB942 | TCCCCCCGGGGTGCGGCCTGAAAAACAGTGC |
| oKB1212 | R ArcZ HindIII | PCR: pKB942 | CCGGCCAAGCTTAAAAAATGACCCCGG CTAGACC |
| oKB1194 | F CyaR XmaI | PCR: pKB911 | TCCCCCCGGGGCTGAAAAACATAACCCATAAA ATGCTAGC |
| oKB1195 | R CyaR HindIII | PCR: pKB911 | CCGGCCAAGCTTAAAAAATAAGCCCGT GTAAGGGAGATTAC |
| oKB1109 | F MgrR XmaI | PCR: pKB856 | TCCCCCCGGGGATTCGTTATCAGTGCAGGAAA ATGCC |
| oKB1110 | R MgrR HindIII | PCR: pKB856 | CCGGCCAAGCTTAAAAAAAACCGCCAG TAAACCGGC |

**Table S4.** Predicted constitutive promoter strengths in pAdapter constructs generated by site-directed mutagenesis. The -35 and -10 sequences are provided for each plasmid; bold letters represent a match to the $\sigma^{70}$ consensus sequence. Plasmids are arranged in order of decreasing predicted strength as determined using the energy binding matrix determined in Brewster et al. 2012.

| Plasmid | Promoter sequence (-35) | (-10) | Relative Strength |
|---|---|---|---|
| **Consensus** | **TTGACA** | **TATAAT** | 3.7 (strongest) |
| pKB989 fully de-repressed | **TT**T**ACA** | **TATAAT** | 2.7 * |
| pKB989 fully repressed (leaky expression) | | | < 2.7 ** |
| pCW14 | AC**GA**T**A** | **TATA**C**T** | 2.2 |
| pCW15 | CC**GA**T**A** | **TATA**C**T** | 1.9 |
| pCW16 | CC**GA**C**A** | **TATA**C**T** | 1.8 |
| pCW17 | AC**GA**T**A** | **TATA**G**T** | 1.8 |
| pCW18 | CC**GA**C**A** | **TATA**G**T** | 1.3 |
| pCW19 | CC**GA**T**A** | G**ATA**G**T** | 1.2 (weakest) |

* pKB989 encodes the CI-MS2$^{CP}$ fusion protein under the control of an IPTG-inducible *lacUV5* promoter. The value listed above is the predicted strength of the fully de-repressed promoter in the absence of any LacI repression.

** When pKB989 is fully repressed (in the absence of IPTG), there is some leaky expression, but this cannot be predicted from the energy binding matrix.

**Table S5.** Dissociation constants for Hfq-sRNA interactions for Hfq mutants from literature.

| | Dissociation Constant ($K_D$; nM) | | | | |
|---|---|---|---|---|---|
| **Hfq variant** | **$A_{27}$[1]** | **DsrA[1]** | **OxyS[2]** | **$A_{27}$[2]** | **DsrA[2]** |
| WT | 39 ± 1 | 32 ± 1 | 1.7 ± 1.1 | .87 ± .35 | .74 ± .4 |
| D9A | 36 ± 1 | 14 ± 1 | | | |
| Y25D | 224 ± 2 | 15 ± 1 | 3.5 ± 1.6 | > 250 | 1.4 ± .13 |
| I30D | 363 ± 3 | 26 ± 1 | 5.9 ± 2.3 | > 250 | 2.1 ± 1.2 |
| Q41A | 26 ± 1 | 25 ± 1 | | | |
| Y55A | 115 ± 2 | 82 ± 4 | | | |
| Y55W | 69 ± 1 | 47 ± 1 | | | |
| K56A | 59 ± 1 | 101 ± 2 | 100 ± 20 | 6.8 ± 3.1 | 180 ± 62 |
| I30D/Y55W | 312 ± 9 | 41 ± 12 | | | |
| Y25D/Y55A | 270 ± 12 | 194 ± 9 | | | |

[1] $K_D$ values measured by gel-shift assays and reported in (Mikulecky et al. 2004)
[2] $K_D$ values measured by filter-binding assays and reported in (Olejniczak 2011).
